# A RIF1/KAP1-based toggle switch stabilises the identities of the inactive and active X chromosomes during X inactivation

**DOI:** 10.1101/2020.06.04.133512

**Authors:** Elin Enervald, Lynn Marie Powell, Lora Boteva, Rossana Foti, Nerea Blanes Ruiz, Gözde Kibar, Agnieszka Piszczek, Fatima Cavaleri, Martin Vingron, Andrea Cerase, Sara B.C. Buonomo

## Abstract

The onset of random X inactivation in mouse requires the switch from a symmetric to an asymmetric state, where the identities of the future inactive and active X chromosomes are assigned. Here we show that RIF1 and KAP1 are two fundamental factors for the definition of the asymmetry. Our data show that at the onset of mESC differentiation, upregulation of the long non-coding RNA Tsix weakens the symmetric RIF1 association with the *Xist* promoter, and opens a window of opportunity for a more stable association of KAP1. KAP1 is required to sustain high levels of Tsix, thus reinforcing and propagating the asymmetry, and, as a result, marking the future active X chromosome. Furthermore, we show that RIF1 association with the future inactive X chromosome is essential for Xist upregulation. This double-bookmarking system, based on the mutually exclusive relationships of Tsix and RIF1, and RIF1 and KAP1, thus coordinates the identification of the inactive and active X chromosomes and initiates a self-sustaining loop that transforms an initially stochastic event into a stably inherited asymmetric X chromosome state.

## INTRODUCTION

X chromosome inactivation (XCI) is the process leading to the stable transcriptional silencing of one of the two X chromosomes in female placental mammals, with the aim of equalising the expression of X-linked genes between males and females (Lyon, 1961). This process represents one of the best-studied examples of how different nuclear processes, such as epigenetic control, 3D organisation of chromatin contacts, sub-nuclear positioning and, potentially, replication-timing regulation, are integrated to achieve transcriptional control. Random XCI (rXCI) is initiated when Xist, an X-encoded long non-coding RNA (lncRNA) is upregulated from one of the two X chromosomes, the future inactive X chromosome (Xi) (Brockdorff, Ashworth et al., 1991, Brown, Ballabio et al., 1991, Marahrens, Panning et al., 1997, Penny, Kay et al., 1996). *In vivo*, this happens around the time of embryo implantation (Monk & Harper, 1978, Rastan, 1982), while in cultured female mouse embryonic stem cells (mESCs), XCI takes place during a narrow time-window at the onset of differentiation (Wutz & Jaenisch, 2000). Monoallelic upregulation of *Xist* is coupled to loss of pluripotency and several activating and repressing factors of this process have been described (Chureau, Chantalat et al., 2011, Furlan, Gutierrez Hernandez et al., 2018, Gontan, Achame et al., 2012, Jonkers, Barakat et al., 2009, Lee & Lu, 1999, Makhlouf, Ouimette et al., 2014, Navarro, Chambers et al., 2008, Tian, Sun et al., 2010). Guided by the three-dimensional (3D) conformation of the X chromosome (Engreitz, Pandya-Jones et al., 2013, Simon, Pinter et al., 2013), Xist spreads *in cis* and recruits SPEN to enhancers and promoters of the X-linked genes to trigger their silencing (Chu, Zhang et al., 2015, Dossin, Pinheiro et al., 2020, McHugh, Chen et al., 2015, Moindrot, Cerase et al., 2015, Monfort, Di Minin et al., 2015) and the exclusion of RNA polymerase II (Chaumeil, Le Baccon et al., 2006, Kucera, Reddy et al., 2011). This in turn promotes the recruitment of the Polycomb Repressive Complexes (PRC1/2) and the accumulation of tri-methylated of H3K27 (H3K27me3) (Sun, Deaton et al., 2006, Zhao, Sun et al., 2008) and monoubiquitinated H2AK119 (H2AK119ub) (de Napoles, Mermoud et al., 2004) on the future inactive X chromosome (Xi). Contextually, Lamin B receptor (LBR) tethers the future Xi to the nuclear periphery to facilitate Xist spreading into active gene regions and the maintenance of the silent state (Chen, Blanco et al., 2016). Finally, the entire Xi switches the replication timing to mid-late S-phase (Takagi, Sugawara et al., 1982). The combination of all these events facilitates the attainment of an exceptionally stable transcriptionally silent status, so robustly controlled that it is maintained throughout the entire life of the organism. One of the least understood of all these steps is the mechanism that, in the initiating phase of XCI, directs the random choice of which one of the two *Xist* alleles to upregulate, marking the future Xi, and which to silence (marking the future active X chromosome, Xa). We will refer to this process as the “choice” (Avner & Heard, 2001). This is a key stage, as failure to establish monoallelic *Xist* expression can result in either both X chromosomes being silenced or both remaining active, consequently leading to embryonic lethality (Borensztein, Syx et al., 2017, Marahrens et al., 1997, Takagi & Abe, 1990). Tsix is a lncRNA encoded by a gene that overlaps, in the antisense orientation, with *Xist* and plays a well-established role as an *in cis* repressor of *Xist* (Lee & Lu, 1999). In female mESCs, *Tsix* is bi-allelically expressed, to become downregulated on one of the two X chromosomes, the future Xi, at the onset of differentiation, hence allowing for *in cis Xist* upregulation (Lee, Davidow et al., 1999, Stavropoulos, Lu et al., 2001). The switch to mono-allelic expression of *Tsix* is important in assigning the destinies of the future Xi (Tsix silenced) and Xa (Tsix transiently maintained). In fact, interfering with the expression of one of the two *Tsix* alleles in female mESCs results in a non-random choice, with the *Tsix*-defective chromosome predetermined as the future Xi (reviewed in (Starmer & Magnuson, 2009)). Although *Tsix* downregulation is essential for *in cis* upregulation of Xist, the molecular mechanism of Tsix-driven silencing is still unclear. The *Tsix* terminator region overlaps with the *Xist* promoter, and *Tsix* transcription through this region and/or Tsix RNA have been proposed to be important for *Xist* repression (Shibata & Lee, 2004) by promoting a silenced chromatin state (Navarro, Page et al., 2006, Navarro, Pichard et al., 2005, Ohhata, Hoki et al., 2008, Sado, Hoki et al., 2005). The establishment of the opposite *Xist/Tsix* expression patterns on the two genetically identical X chromosomes, must rely on the coordinated asymmetric distribution of activators and/or repressors of transcription.

Mathematical modelling can recapitulate the experimental features of XCI by postulating the existence of an *in cis* negative regulator of *Xist* (cXR) and an *in trans*, X-linked, *Xist* activator (tXA) (Mutzel, Okamoto et al., 2019). While a cXR is sufficient to explain the maintenance of mono-allelic *Xist* expression, a tXA is needed to explain 1. the establishment of the *Xist* mono-allelic expression; 2. the female specificity of XCI; 3. the resolution of bi-allelic *Xist* expression detected, to various extents, in different organisms (Mutzel et al., 2019). In mouse, Tsix is the most likely cXR, while RNF12, an X-linked ubiquitin ligase that functions as a dose-dependent trigger of XCI (Gontan et al., 2012, Jonkers et al., 2009), has been proposed as a tXA. However, while over-expression of *Rlim* (*Rnf12*) in male cells can induce XCI (Jonkers et al., 2009), its deletion in females is not sufficient to prevent *Xist* upregulation (Shin, Wallingford et al., 2014, Wang, McCannell et al., 2017). Thus, RNF12 could account for the X-linked aspects of the tXA function, such as female specificity and resolution of bi-allelic expression, but one or multiple other transactivators must be contributing to the control of *Xist*. Moreover, conceptually, the expression level of a single, X-linked gene, does not constitute a switch robust or sensitive enough to be the only element to control a clear-cut bi-stable state for *Xist* (active on one and silent on the other allele). The establishment of *in cis*, self-reinforcing and mutually exclusive circuits on the two *Xist* alleles could create the ultrasensitivity required to generate a binary switch-type of control (Mutzel & Schulz, 2020). Key to this model, is the idea that the initial stochastic events will trigger a chain of local, mutually exclusive and self-sustaining events to bookmark both Xi and Xa.

RIF1 is a multifaceted protein, required for the regulation of several of the nuclear processes that take place during XCI. RIF1 is the only known genome-wide regulator of replication timing (Cornacchia, Dileep et al., 2012, Foti, Gnan et al., 2016, Hayano, Kanoh et al., 2012, Hiraga, Alvino et al., 2014, Peace, Ter-Zakarian et al., 2014, Seller & O’Farrell, 2018, Yamazaki, Ishii et al., 2012). It confines long-range chromatin contacts within the respective boundaries of the nuclear A/B compartments (Gnan S., 2020) and plays an as yet unclear function in the control of gene expression (Daxinger, Harten et al., 2013, Foti et al., 2016, Li, Wang et al., 2017, Tanaka, Muto et al., 2016, Toteva, Mason et al., 2017, Zofall, Smith et al., 2016). RIF1 is an adaptor for Protein Phosphatase 1 (PP1), one of the main Ser/Thr phosphatases in eukaryotic cells (Alver, Chadha et al., 2017, Dave, Cooley et al., 2014, Hiraga et al., 2014, Hiraga, Ly et al., 2017, Mattarocci, Shyian et al., 2014, Sreesankar, Bharathi et al., 2015, Trinkle-Mulcahy, Andersen et al., 2006). In *Drosophila melanogaster*, the RIF1-PP1 interaction was shown to be essential during embryonic development (Seller & O’Farrell, 2018). In addition, the RIF1-PP1 interaction is essential for RIF1-dependent organisation of chromatin contacts (Gnan S., 2020). Others (Chapman, Barral et al., 2013, Daxinger et al., 2013) and we (this work) have observed that mouse RIF1 deficiency is associated with a sex-linked differential lethality, with the female embryos dying around the time of implantation. These data have suggested that RIF1 could be important during XCI. Here we find that RIF1, present biallelically on the *Xist* P2-promoter in female mESCs (Chapman, Cotton et al., 2014, Navarro et al., 2005), becomes asymmetrically enriched at P2 on the future Xi, concomitant with the choice, at the time when *Tsix* expression switches from bi-to mono-allelic. On the future Xi, RIF1 then plays an essential role in *Xist* upregulation. RIF1 removal allows the binding of KAP1 (KRAB-associated protein 1, also known as TRIM28 and TIF1β), regulating Tsix levels at the onset of XCI and influencing to the choice of the future inactive/active X chromosome (Xi/Xa). Our data identify the asymmetric binding of RIF1, and, consequently, of KAP1, as a key steps in the molecular cascade that leads to the bookmarking of the future Xi and Xa, thus implementing a stable binary state for Xist.

## RESULTS

### RIF1 is required for X inactivation during embryonic development and for *Xist* upregulation

The analysis of the progeny derived from inter-crosses of mice heterozygous for a *Rif1* null allele (*Rif1*^*+/-*^, Appendix Fig. S1A and B) in a pure C57BL/6J genetic background has revealed that *Rif1* is essential for embryonic development (Fig. 1A). In contrast, in a mixed genetic C57BL/6J-129/SvJ background, *Rif1* deletion results in a differential lethality between the sexes (Fig. 1B). Indeed, in this case, only a small proportion of the expected *Rif1*^*-/-*^ mice, exclusively males, are recovered at weaning. In order to pinpoint more precisely the time of the onset of lethality, we have analysed the viability of *Rif1*^*-/-*^ embryos at different stages of development in a C57BL/6J pure background. We found that, up to the blastocyst stage, there are no obvious differences between male (our unpublished observation) and female *Rif1* null and wild type embryos (Fig. EV1A). However, immediately after implantation (E5.5/E6.5), the female embryos show a dramatically reduced expansion of the epiblast (Fig. EV1B, C and D) and are already dead by E7.5 (Fig. 1C and D). In contrast, males appear to die only around mid-gestation (Fig. 1C). This early-onset lethality observed specifically in females suggests that the lack of RIF1 could interfere with the process of XCI, as the timing coincides with the onset of random XCI.

**Figure 1.**
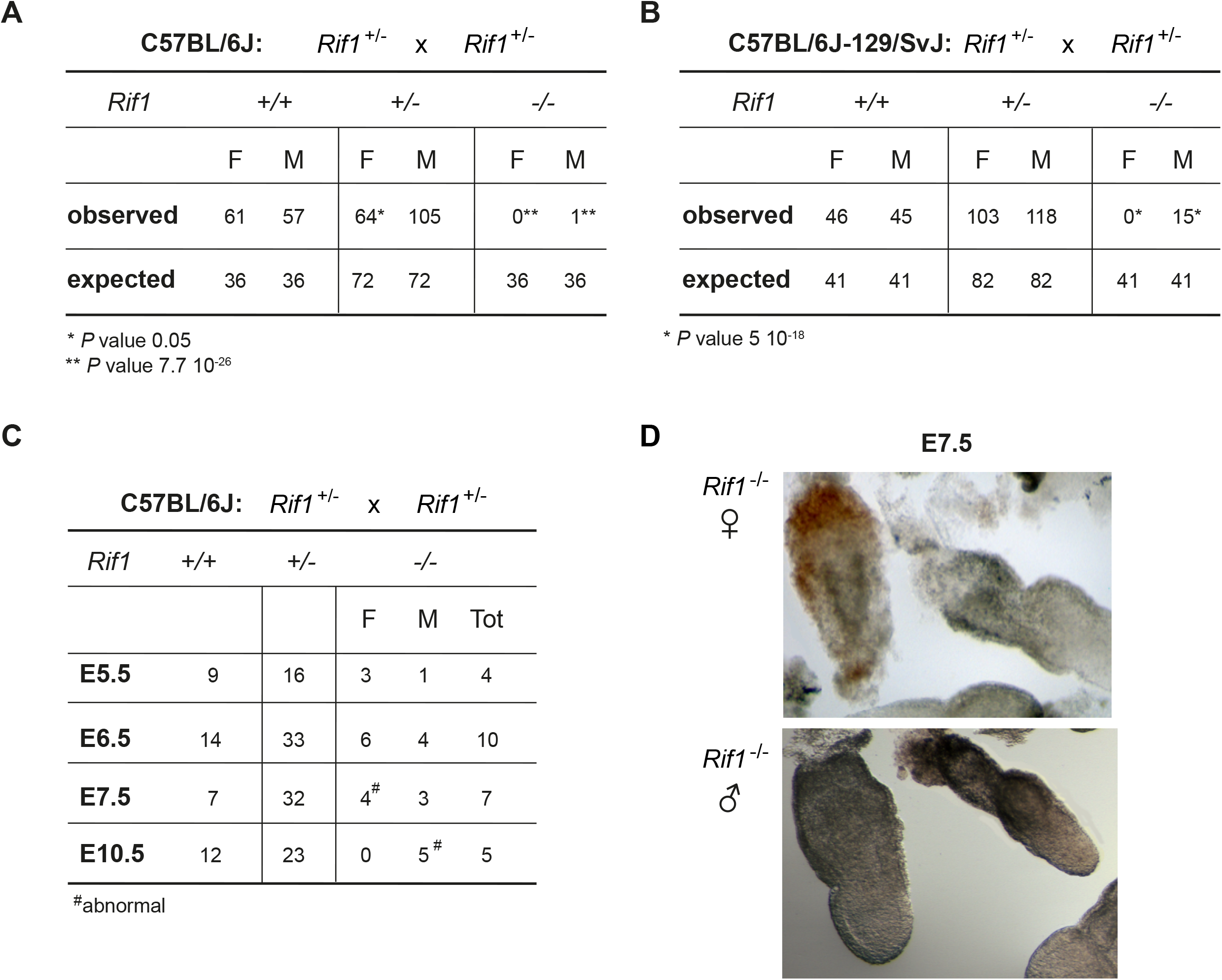
*Rif1* deficiency leads to female embryonic lethality at peri-implantation. Tables summarising the number and the sex of the pups recovered at weaning from *Rif1*^*+/-*^ x *Rif1*^*+/-*^ mice inter-crosses, either in a C57BL/6J **(A)** or in a mixed C57BL/6J-129/SvJ genetic background **(B)**. The observed number of mice is compared to the expected number, based on the Mendelian ratio. *p* calculated by 𝒳^2^. **(C)**. The table summarises the number and the sex of the embryos of the indicated genotypes, recovered from timed matings of *Rif1*^*+/-*^ x *Rif1*^*+/-*^ mice, in a C57BL/6J genetic background. The day of gestation (E) is indicated (**D)**. Representative images of *Rif1*^*-/-*^ E7.5 embryos, female top and male bottom.

Given the diversity of its roles, RIF1 could act at one or several of the multiple steps during XCI. To dissect at what stage(s) of the process RIF1 is required, we generated female mESCs carrying homozygous conditional *Rif1* allele (*Rif1*^*Flox/Flox*^) and a tamoxifen-inducible CRE recombinase (*Rosa26*^*Cre-ERT/+*^,(Buonomo, Wu et al., 2009)). To trigger XCI in the absence of *Rif1*, we have set up a protocol in which we combined differentiation by embryoid bodies (EB) formation (Doetschman, Eistetter et al., 1985) and tamoxifen treatment (Fig. 2A and Materials and Methods). By RT-qPCR as well as by RNA sequencing, we found that *Rif1* deletion (Fig. 2B) severely impairs *Xist* upregulation (Fig. 2C, Fig. EV2A, EV2B and EV3A) and, consequently, the enrichment of H3K27me3 on the future Xi (Fig. 2D). Failure of *Xist* upregulation in the absence of *Rif1* is not due to a general defect in exit from pluripotency (Fig. EV2C and EV3B) or to failed commitment to differentiation (Fig. EV2C and EV3C). Moreover, during the early stages of differentiation the levels of the main negative regulator of Xist, Tsix, appear to be reduced faster in *Rif1* knockout cells compared to the control (Appendix Fig. S2A). Finally, the overall dynamics of RNF12 appear comparable between control and *Rif1* knock out cells (Fig. EV2B and Appendix Fig. S2B). Overall, these results indicate that failure of *Xist* upregulation is the likely cause of defective XCI in *Rif1* null female embryos and that RIF1 could directly and positively regulate *Xist* expression.

**Figure 2.**
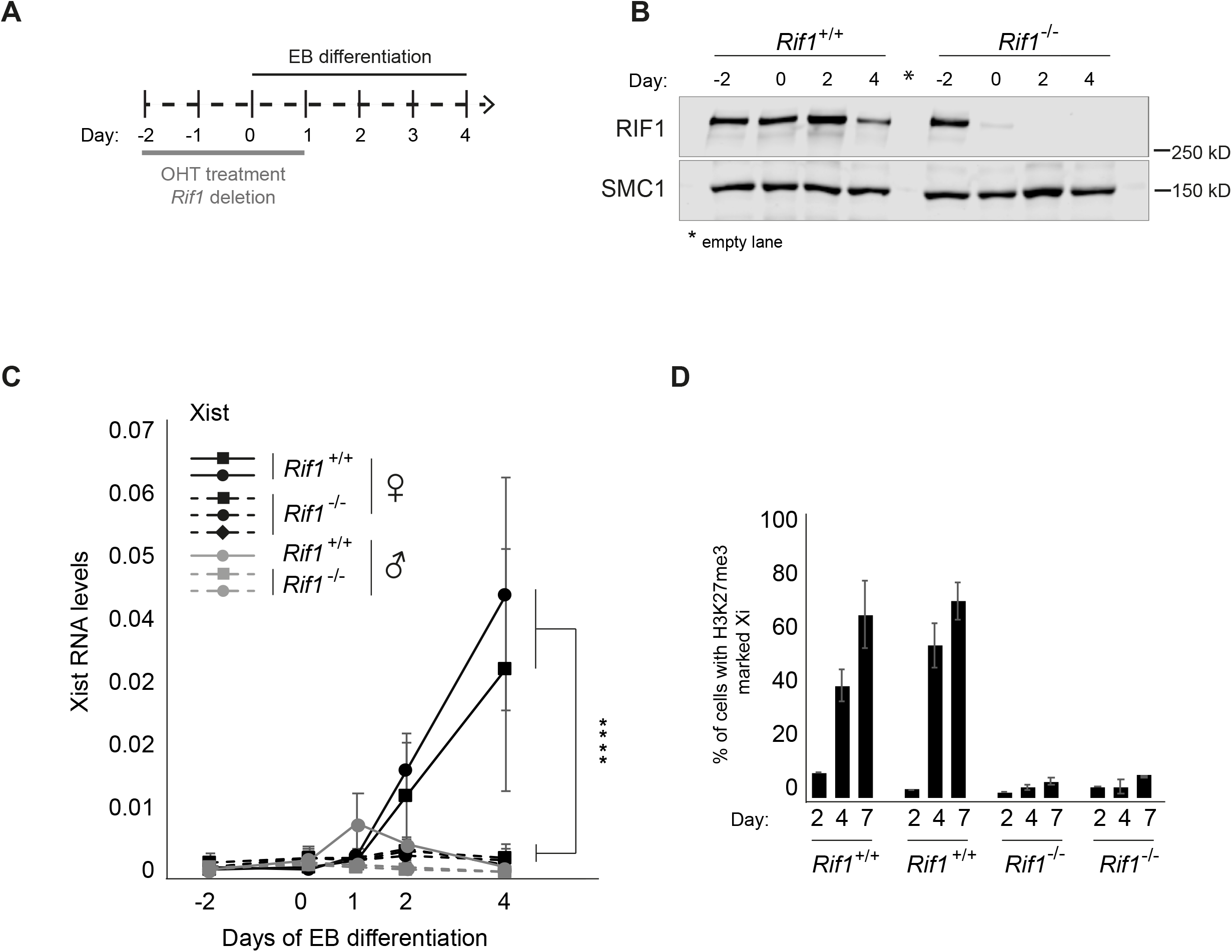
*Rif1* null female mESCs fail to upregulate *Xist* upon differentiation. **(A)**. Overview of the experimental design. *Rif1*^*+/+*^ and *Rif1*^*F/F*^ mESCs were grown for two days in medium supplemented with 4-Hydroxytamoxifen (OHT) to induce the translocation into the nucleus of the Cre-recombinase, leading to *Rif1* deletion in the Rif1^F/F^ cells (*Rif1*^*-/-*^). The embryoid body (EB) differentiation protocol was then started to trigger XCI. OHT was kept in the medium during the first 24 hours of EB differentiation. Cells were differentiated up to 4 (RNA analysis) or 7 days (H3K27me3 IF). **(B)**. Representative western blot to monitor RIF1 levels after Cre-mediated *Rif1* deletion and EB differentiation. SMC1: loading control. **(C)**. Time course analysis of Xist RNA expression by RT-qPCR during EB differentiation of *Rif1*^*+/+*^ (*Rif1*^*+/+*^ +OHT) and *Rif1*^*-/-*^ (*Rif1*^*F/F*^ +OHT) cells at the indicated timepoints. *Rif1*^*+/+*^ (solid line) and *Rif1*^*-/-*^ (dashed line), female (black) and male (grey). Data are presented as mean ± standard deviation from three (female lines) or two (male lines) independent experiments. Statistical significance was determined using 2-way ANOVA comparing female *Rif1*^*+/+*^ to female *Rif1*^*-/-*^ cell lines (****p ≤ 0.0001). Xist RT-primers Xist ex7 F and R were used. Values are normalised to a geometric mean consisting of the expression of *Gapdh, Ubiquitin* and *β-Actin*. **(D)**. Bar plot summarising the number of cells showing H3K27me3-marked Xi as a percentage of total cells counted, in *Rif1*^*+/+*^ (*Rif1*^*+/+*^ +OHT) and *Rif1*^*-/-*^ (*Rif1*^*F/F*^ +OHT) female mESCs at the indicated days of EB differentiation. Averages ±standard deviation from three (day 4 and 7) and two (day 2) independent experiments (n>200).

### RIF1 is a positive regulator of *Xist* and its binding specifically bookmarks the future Xi

*Xist* is controlled from two promoters, P1 and P2 (Johnston, Nesterova et al., 1998), separated by a repetitive region essential for the silencing properties of Xist (Wutz, Rasmussen et al., 2002). While the upstream P1 promoter gives rise to the Xist main transcript, P2 appears to serve as an internal regulatory unit, possibly controlling P1 expression (Makhlouf et al., 2014). We found that RIF1 is enriched specifically at *Xist* P2 promoter, both in mESCs (Fig. 3A, Appendix Fig. S2C and S2D) and in early EBs (Fig. 3B), supporting the hypothesis that RIF1 could be a direct regulator of *Xist* expression. In agreement with this, we found that P2 harbours two potential RIF1-binding sites, defined by the presence of a consensus derived from the analysis of RIF1 genome-wide distribution by ChIP-seq in female mESCs (Foti et al., 2016) (Appendix Fig. S3A). To confirm that RIF1 association with *Xist* promoter has a positive effect on *Xist* expression, we used a reporter assay system, where *Xist* promoter has been cloned upstream of a firefly Luciferase gene (Gontan et al., 2012). We found that, upon differentiation, in the absence of RIF1, the induction of Luciferase from the *Xist* promoter is significantly reduced (Fig. 3C), supporting the hypothesis that RIF1 association with P2 exerts a positive, direct effect on *Xist* transcription.

**Figure 3.**
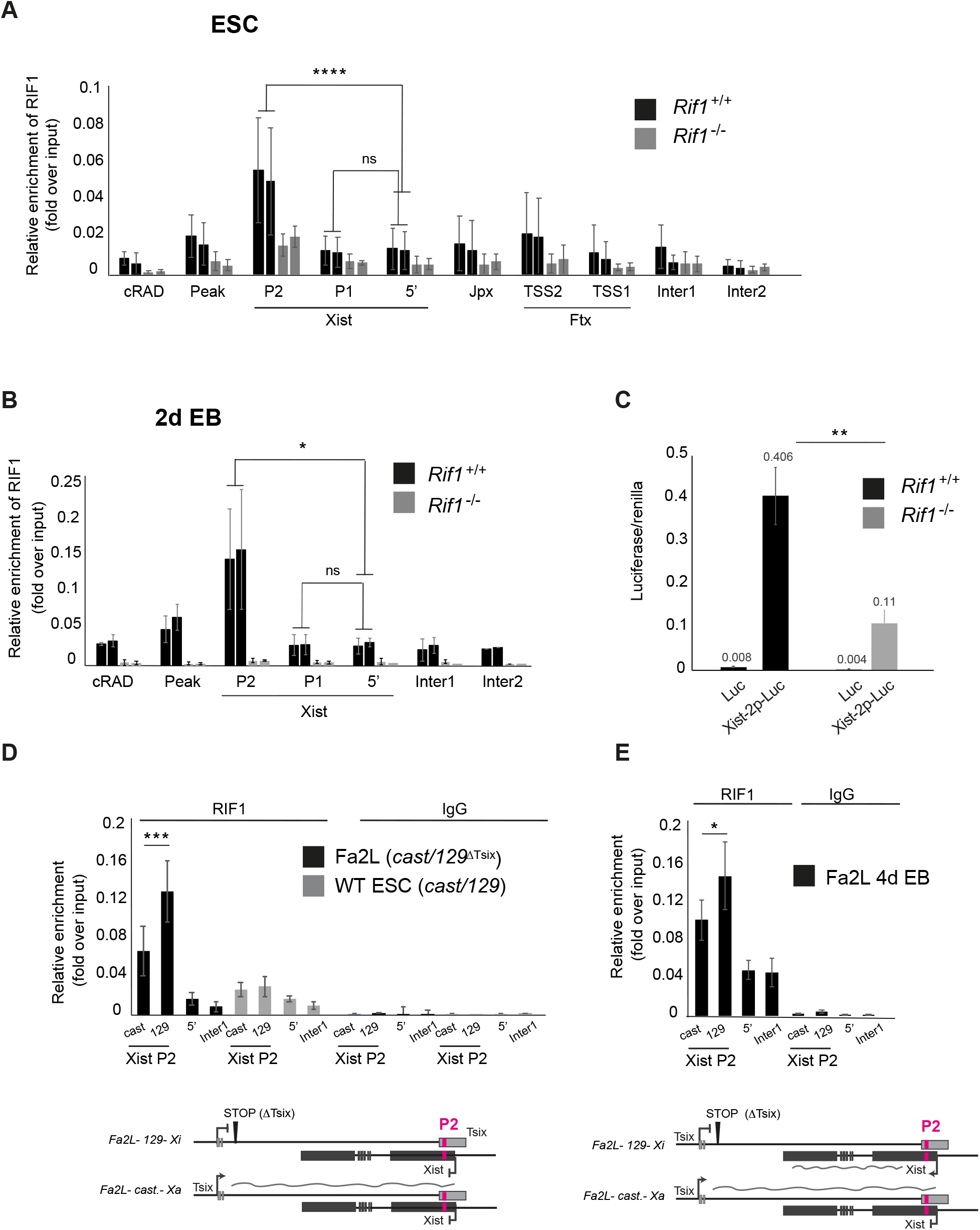
RIF1 associates with *Xist* promoter on the future Xi. RIF1 association with the *Xist* promoter assessed by ChIP-qPCR in two independent *Rif1*^*+/+*^ (*Rif1*^*+/+*^ +OHT, black) and two *Rif1*^*-/-*^ (*Rif1*^*F/F*^ +OHT, grey) female cell lines, in ESCs **(A)**. and at 2 days of EB differentiation **(B)**. P1 and P2 indicate the two *Xist* promoters, 5’ indicates a region 2 kb upstream of *Xist* TSS. Inter1 and 2 are two intergenic regions that serve as negative controls. Peak and cRAD represent two previously identified regions of RIF1 association (positive control). See Appendix Fig. S2C for primer positions within Xist. Mean ±standard deviation from 3 independent experiments (A) and 2 independent experiments (B). *p* calculated by Student’s two-tailed, paired *t* test comparing RIF1 association in *Rif1*^*+/+*^ cells on Xist P2 and P1 versus 5’. **(C)**. *Rif1* deletion decreases the efficiency of upregulation of a *Luciferase* reporter under the control of *Xist* promoter (Xist-2p-luc), at 2 days of EBs differentiation. As a control (Luc), empty luciferase reporter vector, was transfected in parallel. The average of three independent experiments is shown. Error bars indicate the standard deviation. *p* calculated by Student’s two-tailed, unpaired *t* test, for comparison of fold activation of Xist-2p-Luc normalised to empty vector (Luc) in *Rif1*^*+/+*^ versus *Rif1*^*-/-*^ cells. **p ≤ 0.01. See Supplementary Methods for details about the normalisation. **(D)**. Association of RIF1 with *Xist* P2 in the Fa2L cells (black) and a wild type female mESC line (grey), also harbouring one *castaneus* and one *129* X chromosome. Allele-specific ChIP-qPCR primers were used, *cast* indicates association with the *castaneus Xist* P2 and *129* indicates association with the *129 Xist* P2. Enrichments are presented relative to input DNA. Mean ±standard deviation from 3 independent experiments. *p* calculated by Student’s two-tailed, paired *t* test comparing RIF1 association with the *castaneus* and with the *129* X chromosome Xist P2. Below, schematic of the *Xist/Tsix* alelles in the Fa2L cells. **(E)** Association of RIF1 with *Xist* P2 in the Fa2L cells (black) upon differentiation. The analysis was performed as in (D). *p ≤ 0.05, ***p ≤ 0.001, ****p ≤ 0.0001 and ns=not significant. Below, schematic of the *Xist/Tsix* alelles in the Fa2L differentiated cells.

Upon differentiation, *Xist* is mono-allelically transcribed, upregulated only from the future Xi. If RIF1 acts as a positive regulator of *Xist*, we would expect it to be associated mono-allelically, specifically with P2 on the future Xi. In order to test this hypothesis, we have taken advantage of the Fa2L cell line, in which 1. the two X chromosomes can be discriminated, as one originates from *Mus castaneus* (*cast*) and the other from *Mus musculus 129/SvJ* (*129*) mouse strains; 2. Xa (*cast*) and Xi (*129*) are predetermined, as the *129 Tsix* allele carries a transcriptional stop signal, approximately 4kb downstream from the *Tsix* major promoter (Fig. 3D, scheme and (Luikenhuis, Wutz et al., 2001)). *Xist* is therefore preferentially upregulated from the *129*-derived X chromosome. We have analysed the association of RIF1 with *Xist* P2 promoter of the future Xa and Xi by allele-specific ChIP-qPCR (Appendix Fig. S3B) and found that RIF1 is preferentially associated with the *Xist* P2 promoter of the*129 Xist* allele (future Xi) in both mESCs (Fig. 3D) and upon differentiation (Fig. 3E). Importantly, in control wild type mESCs (bi-allelically expressed *Tsix*), also carrying one *cast* and one *129* X chromosome, RIF1 is equally distributed on both P2 promoters (Fig. 3D), suggesting that the asymmetric association with the future Xi is concomitant with the switch from bi-to mono-allelic *Tsix* expression that accompanies the choice and allows *Xist* monoallelic upregulation. As in the case of RIF1 conditional cells, depletion of RIF1 in Fa2L cells (Appendix Fig. S3C) also compromises *Xist* upregulation (Appendix Fig. S3D). These data show that RIF1’s asymmetric association with the future Xi parallels the choice and it is essential for *Xist* upregulation.

### KAP1 association with the future Xa is important for the onset of XCI

With the aim of understanding the molecular mechanism by which RIF1 regulates *Xist* expression, we have investigated whether some of the known transcriptional regulators associated with RIF1 are also required for XCI. We focused in particular on KAP1, as KAP1 and RIF1 have already been shown to regulate overlapping targets, such as Dux and MERVLs (Li et al., 2017, Maksakova, Thompson et al., 2013, Percharde, Lin et al., 2018). We found that knock down of Kap1 (Fig. 4A and B) impairs *Xist* upregulation (Fig. 4C and Fig. EV4A), similarly to the knockout of *Rif1*. This is not due to compromised exit from pluripotency (Fig. EV4B), impaired activation of the differentiation transcriptional program (Fig. EV4C) or reduced RIF1 levels (Fig. EV4D), suggesting that diminished *Xist* activation is not a consequence of an overall impaired cell differentiation. In addition, the dynamics of expression of RNF12 appear comparable between control and Kap1 knock down cells (Fig. EV4E).

**Figure 4.**
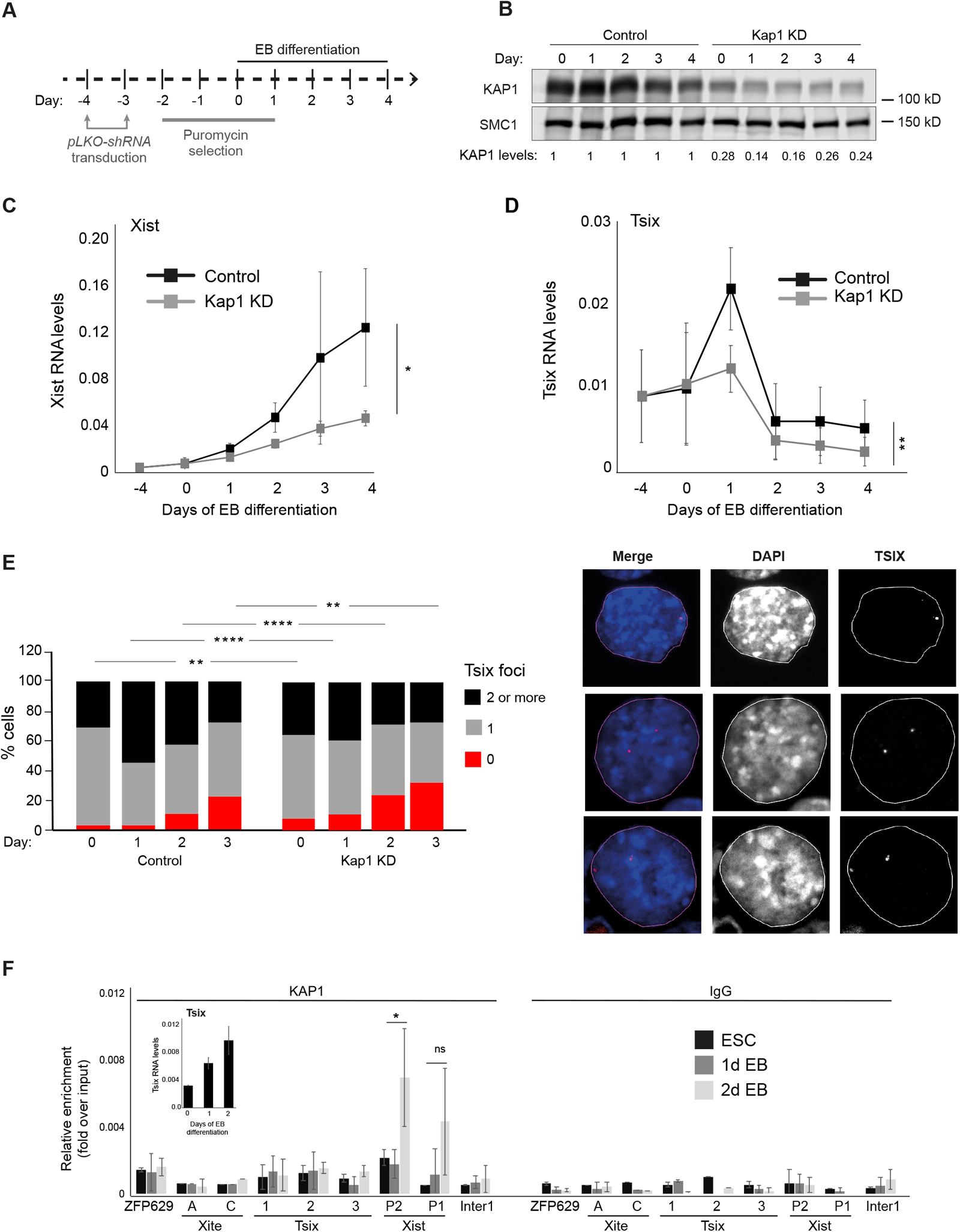
KAP1 associates with *Xist* promoter on the future Xa. **(A)**. Schematic of the experimental design. **(B)**. Western blot analysis of KAP1 levels after knock down, in female mESCs and during EB differentiation. SMC1: loading control. Below, quantification of KAP1 protein levels relative to control cells. **(C**). Time course analysis of *Xist* expression by RT-qPCR during EB differentiation of female mESCs following knock down of Luciferase (Control, black) and Kap1 (Kap1 KD, grey), at the indicated timepoints. Data are presented as mean ±standard deviation from three independent experiments. Statistical significance was determined using 2-way ANOVA. *Xist* primers Xist ex3 F and Xist ex4 R were used. Normalisation was performed using a geometric mean consisting of the expression of *Rplp0, Ubiquitin* and *Sdha*. **(D)**. Tsix RNA levels in female mESCs infected with shRNA directed against Luciferase (Control, black) and KAP1 (Kap1 KD, grey), during differentiation. Mean ±standard deviation from four independent experiments are shown. Statistical significance was determined using 2-way ANOVA. Values have first been normalised to a geometric mean consisting of the expression of *Rplp0, Ubiquitin* and *Sdha*. **(E)**. RNA FISH analysis of *Tsix* expression during differentiation of female mESCs expressing an shRNA directed against Luciferase (control) or against Kap1 (Kap1 KD). Left: Cells with no (0, red), one (1, grey) and two or more (2, black) Tsix foci were counted in two independent experiments, shown averaged. Statistical significance was determined by 𝒳^2^. A minimum of 110 cells were counted per each time point for each line. Right: examples of cells with one (top) or two (central and bottom) Tsix FISH signals. **(F)**. ChIP-qPCR analysis of KAP1 association with the indicated sites in wild type female mESCs (*Rif1*^*+/+*^, same as used in Fig. 3B but without OHT) and during early differentiation. ZFP629 is a well-characterised KAP1 associated region (positive control). Xite A and C indicate two regions within the Tsix enhancer *Xite*, Tsix region 1 indicates Tsix major promoter, Tsix region 2 indicates the Dxpas34 region, Tsix region 3 indicates a region slightly downstream of the Dxpas34 region. P1 and P2 indicate the two *Xist* promoters, 5’ indicates a region 2 kb upstream of *Xist* TSS. Inter1 is an intergenic region. See Appendix Fig. S2C for the positions of the primers within *Xist* and *Tsix*. The data are presented as mean ±standard deviation from three (2d EB and 1d EB) and two (ESCs) independent experiments. Statistical significance was calculated by Student’s two-tailed unpaired *t* test comparing RIF1 association to Xist P2 and P1 in 2d EB versus 1d EB (*p ≤ 0.05, **p ≤ 0.01, ***p ≤ 0.001, ****p ≤ 0.0001 and ns=not significant). In the inset, Tsix RNA levels were quantified by RT-qPCR during the differentiation of wild type female ESCs shown in Fig. 4F. The average of two experiments is shown. Tsix values are normalised to a geometric mean consisting of the expression of *Rplp0, Ubiquitin* and *Sdha*. Error bars indicate standard deviations.

In wild type cells, during the early stages of differentiation, Tsix levels rise transiently (at 1, or 1 and 2 days of embryoid body differentiation respectively, depending on the culture conditions, Fig. 4D and Appendix Fig. S2A). The boost corresponds to an increased detection of Tsix RNA from both alleles (Fig. 4E), suggesting that this step precedes the switch to *Tsix* mono-allelic expression and the consequent choice of Xa/Xi. Upon KAP1 downregulation, we found not only a failure in the temporary boost of Tsix levels (Fig. 4D), but also a failure to evolve towards Tsix mono-allelic expression, as Tsix becomes undetectable (Fig. 4E). Interestingly, in wild type undifferentiated ESCs, KAP1 is barely detectable on *Xist* P2, where it is specifically recruited concomitantly with the boost of Tsix levels at the early stages of differentiation (Fig. 4F). These data suggest that, unlike RIF1, KAP1 could be indirectly required for *Xist* control, by ensuring the dynamic regulation of Tsix levels that leads to/precedes the switch from bi- to mono-allelic and the consequent choice. In support of this hypothesis, we found that in the Fa2L cells, where *Tsix* truncation results in obligatory mono-allelic expression, KAP1 is preferentially associated with P2 on the *castaneus* allele (future Xa), which expresses full-length Tsix both in undifferentiated cells (Fig. 5A), and upon differentiation (Fig. EV4F). Altogether, these data suggest that the failure to upregulate *Xist* caused by *Rif1* deletion and by Kap1 knock down have different causes. While RIF1 is directly required to promote *Xist* upregulation, KAP1’s function is to drive the transient increase of Tsix levels that precedes the choice. The consequent failure to upregulate *Xist* could be caused, in this case, by a failure to execute the choice, evolving the low, bi-allelic Tsix levels typical of ESCs directly towards Tsix absence, rather than asymmetric distribution. If this hypothesis were correct, we would expect that Kap1 knock down in a “forced-choice” situation, like in the Fa2L cells, would have no consequences on *Xist* upregulation. Confirming our hypothesis, we found that Kap1 knock down (Appendix Fig. S4A) does not affect Xist levels in the Fa2L cells (Fig. 5B), with Tsix levels remaining low also upon differentiation (Appendix Fig. S4B).

**Figure 5.**
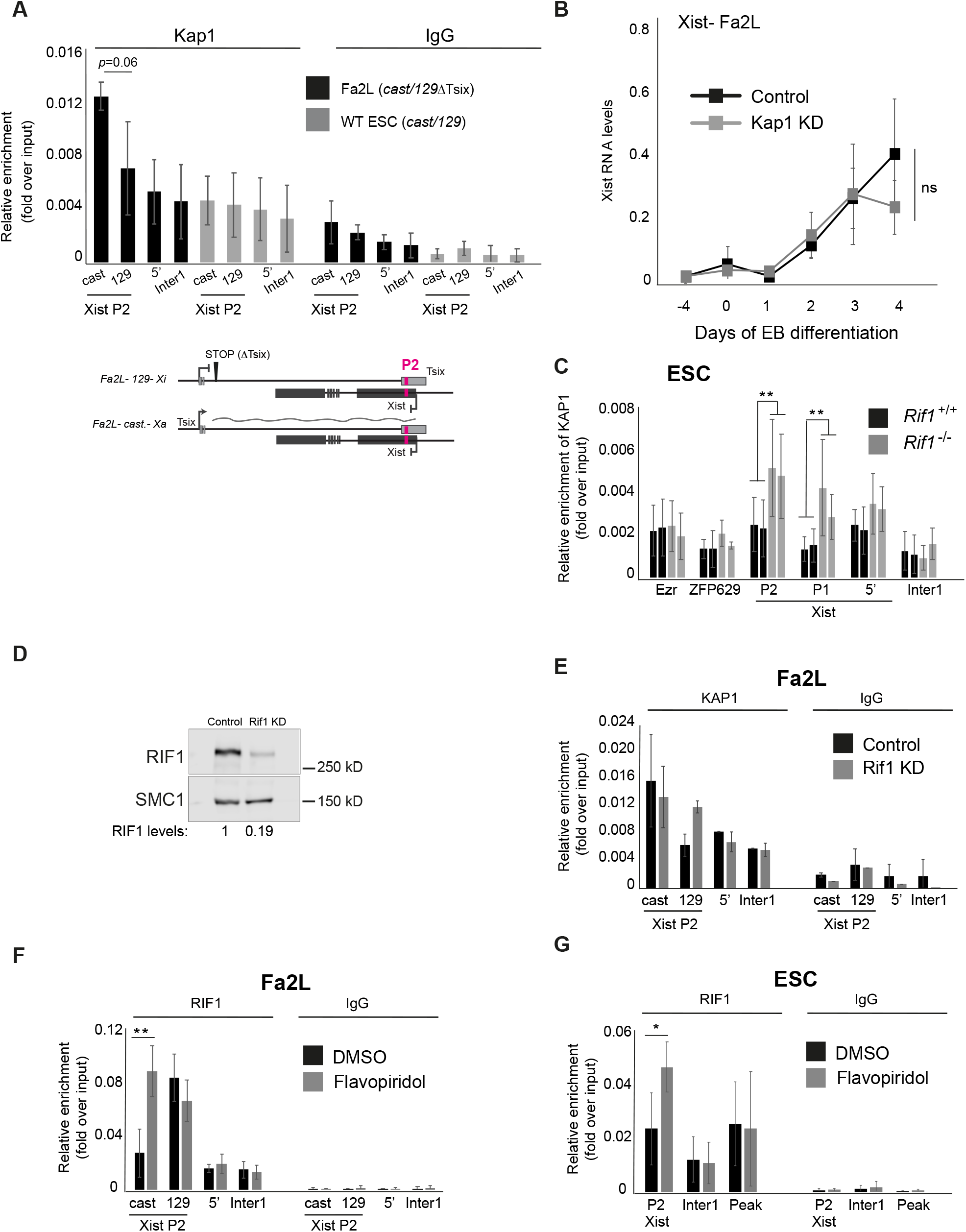
*Tsix* expression controls Rif1 association with Xist P2. **(A)**. Using allele-specific primers, ChIP-qPCR was used to measure the association of KAP1 with *Xist* P2 in the Fa2L cells (black) and a wild type female mESC line also harbouring one *castaneus* and one *129* X chromosome (grey). *cast* indicates association with the *castaneus Xist* P2 and *129* indicates association with the *129 Xist* P2. Enrichments are presented relative to input DNA. Mean ±standard deviation from a minimum of three independent experiments. Statistical significance was determined by Student’s two-tailed, paired *t* test. Below, schematic of the *Xist/Tsix* alleles in the Fa2L cells. **(B)**. RT-qPCR analysis of *Xist* expression levels in the Fa2L cell line, following expression of shRNA against Luciferase (Control, black) and Kap1 (Kap1 KD, grey), at the indicated timepoints. 2-way ANOVA was used to determine statistical significance. ns=not significant. **(C)**. KAP1 association with *Xist* promoter in two independent *Rif1*^*+/+*^ (*Rif1*^*+/+*^ +OHT, black) and two *Rif1*^*-/-*^ (*Rif1*^*F/F*^ +OHT, grey) female mESC cell lines. *Ezr* is an additional region known to be associated with KAP1 in mESCs. Enrichments are presented relative to input DNA. Mean ±standard deviation from a minimum of three independent experiments per cell line are displayed. Statistical significance was determined using Student’s two-tailed, unpaired *t* test comparing the KAP1 association with Xist P2 and P1 in *Rif1*^*+/+*^ versus *Rif1*^*-/-*^ cells. **(D)**. Western blot analysis of RIF1 levels in protein extracts from Fa2L cells after Rif1 knock down. SMC1: loading control. Below, quantification of RIF1 protein levels compared to control cells. **(E)**. Allele-specific KAP1 association with *Xist* P2 in Fa2L cells following knock down of Luciferase (Control, black) and Rif1 (Rif1 KD, grey). *cast* indicates association with the *castaneus Xist* P2 promoter and *129* indicates association with the *129 Xist* P2 promoter. Enrichments are presented relative to input DNA. Average ±standard deviation of two independent experiments **(F)**. Quantification by ChIP-qPCR of RIF1 association with the indicated regions in the Fa2L cell line, following treatment with DMSO only (black) or flavopiridol (grey). Primers as in (E). **(G)**. Same as in **(F)** but for a wild type female mESC line. All enrichments are presented relative to input DNA. Mean ±standard deviation from three **(F)** and two **(G)** independent experiments are presented. Statistical significance was determined using Student’s two-tailed, paired *t* test (*p ≤ 0.05, **p ≤ 0.01 and ns=not significant).

### RIF1 negatively regulates KAP1 association with the Xist promoter in mESCs

Our data show that during the initial stages of differentiation, RIF1 asymmetric association with the future Xi parallels the asymmetric binding of KAP1 on the future Xa, suggesting the existence of a double-bookmarking system to identify the future Xi/Xa. In order to understand if and how these two systems are coordinated, we have investigated whether RIF1 regulates KAP1 association with *Xist* P2. Although normally KAP1 is barely detectable at the *Xist* promoter in wild type mESCs and gains association only upon differentiation (Fig. 4F), we found that *Rif1* deletion leads to earlier KAP1 binding to *Xist* promoter, as it becomes already evident in mESCs (Fig. 5C). This is not due to a general increase of *Kap1* expression (Fig. EV2A), KAP1 protein levels (Fig. EV5A) or its overall binding to chromatin (Fig. EV5B). Moreover, KAP1 enrichment is specific for *Xist* promoter, as other regions known to be associated with KAP1 that we have tested, like *Znf629* (Fig. 5C and our unpublished observation) (Ding, Bergmaier et al., 2018), did not show an increased KAP1 association upon *Rif1* deletion. Importantly, the effect of RIF1 deficiency is unlikely to be due an indirect, general remodelling of the *Xist* promoter chromatin, as the association of another P2-specific transcription factor and *Xist* activator, the Yin-Yang-1 (YY1) (Makhlouf et al., 2014), is unchanged in *Rif1* knockout cells (Fig. EV5C). We also found that knocking down Rif1 in undifferentiated Fa2L cells (Fig. 5D) facilitates KAP1 association to *Xist* P2 (Fig. 5E) comparably to what happens in *Rif1* conditional cells upon induction of *Rif1* deletion (Fig. 5C). In the case of the Fa2L cells (Fig. 5E), we could also determine that KAP1 gains access specifically to the future Xi (*129* allele, carrying the truncated *Tsix* allele), where normally RIF1 is preferentially localised (Fig. 3D and E). Overall, these data indicate that, in mESCs, RIF1 is symmetrically associated with *Xist* P2 on both X chromosomes, protecting P2 from the binding of KAP1. Upon triggering differentiation, the transition of RIF1 to an asymmetric association with *Xist* promoter on the future Xi, allows the upregulation of *Xist in cis* and the association of KAP1 with *Xist* promoter on the future Xa, which in turn sustains the increase of Tsix levels that precede the switch to *Tsix* mono-allelic expression and the choice.

### RIF1 asymmetric localisation on the future Xi is driven by *Tsix* expression

How is the transition from bi- to mono-allelic RIF1 association with *Xist* promoter regulated? While this is generally triggered by differentiation, in undifferentiated Fa2L cells it is pre-determined and RIF1 is preferentially associated with the X chromosome that does not express full-length Tsix transcript (Fig. 3D and E). This suggests that Tsix RNA and/or transcription could destabilise RIF1 association with *Xist* promoter. In agreement with this hypothesis, we found that blocking *Tsix* expression by treating mESCs with the CDK9-inhibitor flavopiridol (Chao & Price, 2001) (Fig. EV5D) or, briefly, with triptolide, an inhibitor of transcription initiation (Fig. EV5E), is sufficient to revert RIF1 preferential association with the future Xi in the Fa2L cells to a symmetric mode of binding (Fig. 5F and EV5F). In addition, flavopiridol treatment of wild type mESCs also leads to an increased P2-association of RIF1 (Fig. 5G), indicating that this is not an effect specific to the Fa2L cells. Finally, while this work was under review, RIF1 has been found associated with Tsix RNA in mESCs (Aeby, Lee et al., 2020), supporting the hypothesis that Tsix RNA can compete for RIF1 association with *Xist* P2 in the genome.

## DISCUSSION

While marsupials have adopted an imprinted X inactivation strategy, eutherians have evolved a mechanism based on the random choice of the X chromosome to be inactivated. The latter can contribute to a higher degree of resistance of females to pathogenic X-linked mutations and increase phenotypic diversity. Despite its importance, the mechanisms guiding the random choice are still unclear, partially because of the randomness and consequent heterogeneity in the cell population, partially because of the inaccessibility of the early embryos, where the process takes place naturally and, finally, because of the inherent difficulty of identifying asymmetry involving two identical chromosomes.

Several lines of evidence suggest that *Tsix* is involved in the choice-making process. For example, introduction of a stop codon that blocks Tsix transcript before its overlap with *Xist* (Luikenhuis et al., 2001), or deletions of its major promoter (Vigneau, Augui et al., 2006), or of the GC-rich repeat region that immediately follows it (Dxpas34) (Lee & Lu, 1999), or insertion of a gene trap in the same region, that abolishes the production of Tsix RNA (Sado, Wang et al., 2001), result in a non-random choice, with the *Tsix*-defective chromosome as the future Xi. Moreover, monoallelic downregulation of Tsix levels by deleting *Xite*, a *cis*-acting element that positively regulates *Tsix*, also skews the choice (Ogawa & Lee, 2003). Interestingly, Xist itself can influence the choice, in a yet-to-be-understood feedback control loop. *Xist* ectopic upregulation can in fact skew the choice in favour of the *Xist*-overexpressing chromosome (Nesterova, Johnston et al., 2003, Newall, Duthie et al., 2001).

Our experiments show that RIF1 association with the *Xist* P2 promoter is negatively regulated by *Tsix* expression or RNA levels. Tsix could therefore be the determinant of the asymmetric association of RIF1 with the future Xi at the choice. We would like to propose a model (Fig. 6) whereby, at the onset of differentiation, the transient, biallelic increase of Tsix levels will promote a weaker or more dynamic association of RIF1 with *Xist* P2, thus creating a window of opportunity for KAP1 stochastic association with either allele. The KAP1-bound allele will go on to sustain higher Tsix steady state levels *in cis*, thus skewing RIF1 association with the opposite allele, and initiating a self-reinforcing loop on the future Xa. On the future Xi, RIF1 will promote Xist upregulation, thus establishing the inactivation. The negative effect of RIF1 on KAP1 association with *Xist* promoter in ESCs is at the heart of the mutual exclusion, reinforced by KAP1’s positive effect on the levels of Tsix, that is, in turn, a negative regulator of RIF1 association with *Xist* promoter. How RIF1 excludes KAP1 is currently unclear, but we can envisage at least two potential mechanisms, based either on RIF1/KAP1 competition for binding to a shared site or protein partner, or through KAP1 de-phosphorylation by RIF1-associated PP1. Phosphorylation of KAP1 has indeed been shown to regulate KAP1 association with heterochromatin protein 1 (HP1) (Chang, Chou et al., 2008).

**Figure 6.**
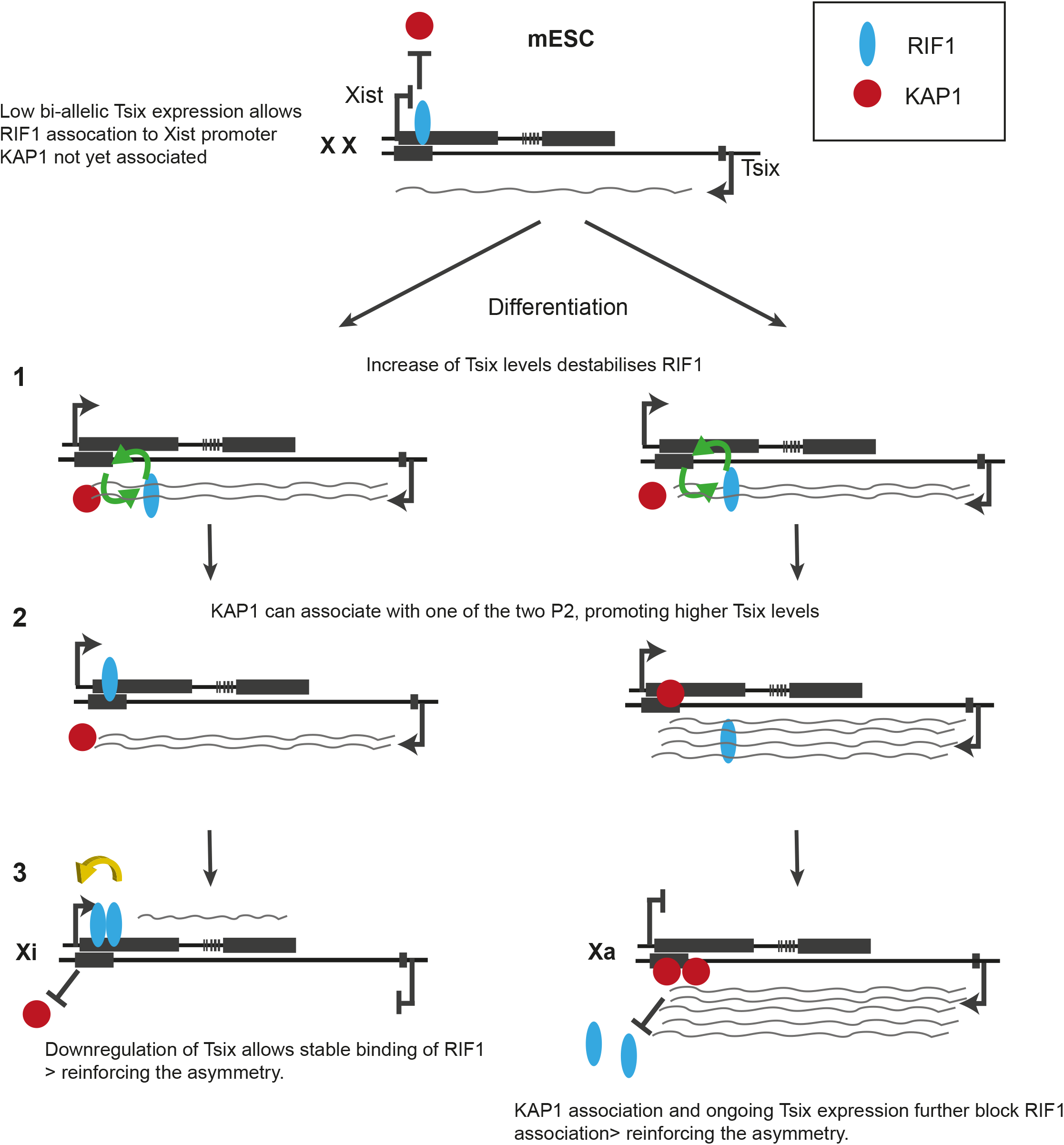
Model for RIF1 and KAP1-dependent bookmarking of Xi and Xa respectively. The low bi-allelic expression of *Tsix* in mESCs allows the association of RIF1 with P2 on both *Xist* alleles. However, the presence of pluripotency-dependent inhibitors will not allow *Xist* upregulation, despite the presence of RIF1. (1) Upon differentiation, the increase of Tsix levels weakens the association of RIF1 with P2. This opens the opportunity for a stochastic KAP1 binding to P2 of one of the two alleles (2). KAP1 is required for sustained high levels of Tsix, further reinforcing RIF1 exclusion from the KAP1-bound/Tsix high allele, and establishing the asymmetry. It is not known whether KAP1 gains access to P2 to promote the increase of Tsix levels first, or whether the increase of Tsix levels is initially triggered by a differentiation-dependent factor. (3). The pluripotency Xist inhibitors having been silenced, RIF1 promotes *Xist* expression on the future Xi. A self-sustainable binary switch is thus created and it consolidates the choice of the future Xi and Xa.

In support of our model, we have shown that, the association of KAP1 with the P2 region upon differentiation coincides with the detection of higher levels of Tsix RNA (Fig. 4F), and this increase is dependent upon KAP1 (Fig. 4D and E). The molecular mechanism by which KAP1 modulates Tsix levels is currently unknown. KAP1 is part of the 7SK complex (McNamara, Reeder et al., 2016), a ribonucleoprotein complex with roles both at the promoter and in the transcriptional termination of several genes, including several lncRNAs (Bunch, Lawney et al., 2016). The region of KAP1 association with the *Xist* promoter coincides with the *Tsix* terminator in the opposite orientation. It is therefore possible that KAP1, as a part of the 7SK complex, could regulate *in cis* Tsix termination and, consequently, stability (reviewed in (Peck, Hughes et al., 2019)). Alternatively, KAP1/7SK could promote *in cis Tsix* expression during the initial stages of differentiation through a terminator-promoter positive feedback loop (Tan-Wong, French et al., 2008). In support of either of these hypotheses, we have found that, as for KAP1, the 7SK complex is also enriched on *Xist* promoter/*Tsix* terminator in *Rif1* knockout mESCs, and it is associated with the future Xa in Fa2L cells, in a KAP1-dependent manner (our unpublished observation). Finally, we cannot exclude a model where KAP1 promotes Tsix increase *in trans*, through a yet unknown differentiation-induced factor. In this case, the association of KAP1 with *Xist* P2 could still contribute *in cis* to the identification of Xa, by establishing a stable repression of *Xist* promoter, with RIF1 shielding the future Xi by excluding KAP1. However, our data do not support this hypothesis. KAP1’s best known repressive function is mediated through its interaction with SetDB1, an HMTase that tri-methylates K9 on histone H3 (H3K9me3). Conversely, we found that in Fa2L cells, the gain of H3K9me3 on the *Xist* promoter of the Xa is independent of KAP1 (our unpublished observation). Yet, KAP1 association with P2 on the future Xa could promote *Xist* stable repression via a mechanism different from tri-methylation of H3K9. Moreover, in disagreement with the hypothesis of KAP1 acting as an *in cis Xist* repressor, we did not detect bi-allelic *Xist* expression in Kap1 knock down cells (Fig. EV4A).

Our data show that the increase of Tsix that precedes and, possibly, leads to a proficient choice, requires KAP1. It has been previously shown that failure to set up the choice as a consequence of homozygous deletion of *Tsix*, leads to a mixture of cells showing either no *Xist* upregulation or bi-allelic upregulation during differentiation (Lee, 2002, Lee, 2005). This is different from what we observe in Kap1 knock down cells, where we detect defective *Xist* up-regulation, but not bi-allelic expression. Nonetheless, a situation where, from the start of the process in ESCs, Tsix is always absent, as in the case of *Tsix*^*-/-*^, is clearly different from the system where Tsix levels are “normal” until differentiation is triggered, as in the case of Kap1 knock down (Fig. 4D).

The dramatic failure of expansion of the epiblast in *Rif1*^*-/-*^ females and the early embryonic lethality described here, contrast with the milder effect of *Xist* conditional inactivation in the epiblast described previously (Yang, Kirby et al., 2016). However, beside the technical differences between a conditional system, where the efficiency of the deletion can be lower than 100%, and a knockout, RIF1 has at least two other key roles, in the regulation of the replication-timing program (Cornacchia et al., 2012, Foti et al., 2016, Hayano et al., 2012, Yamazaki et al., 2012) and replication fork protection (Buonomo et al., 2009, Garzon, Ursich et al., 2019). In fact, depending on the genetic background, most or some of the male embryos also die, although later during development (this work). We cannot therefore exclude that the early female lethality could derive from a synthetic effect of multiple problems, added on top of the failure of X inactivation.

In summary, we propose that, during the stochastic phase of the choice of the future Xi, the asymmetric distribution of RIF1, following a Tsix-dependent stripping of RIF1 from the future Xa, triggers the establishment of two, mutually exclusive, *in cis* circuits that will identify Xi and Xa. RIF1’s presence on P2, inhibiting KAP1 and promoting *Xist* expression on Xi, and KAP1’s presence on P2, sustaining Tsix levels and, thus, helping to exclude RIF1 from Xa, would evolve the initial stochastic event of Tsix asymmetry into a binary switch, where a bi-stable, self-sustaining situation on the two X chromosomes is propagated.

## MATERIALS AND METHODS

### mESC differentiation

Wild type ESCs were plated onto non-coated petri dishes at a concentration of 1×10^6^ cells/ 10 cm^2^, in a volume of 10 ml medium lacking 2i and LIF. At day 4 of differentiation the aggregated embryoid bodies (EBs) were gently transferred to gelatinised tissue culture dishes. Medium was gently changed every 48 hours with minimal disruption of the EBs. EBs were grown for up to 4 or 7 days in total. In experiments where cell differentiation was combined with *Rif1* deletion, the differentiation was preceded by 48 hours of 4-hydroxytamoxifen (OHT, #H7904, Sigma-Aldrich) treatment, at a concentration of 200 nM in ES medium containing LIF and 2i. Differentiation was then started with 2×10^6^ cells/ 10cm^2^ dish for *Rif1*^*+/+*^ and 2.5×10^6^ cells/ 10cm^2^ for *Rif1*^*F/F*^ cells in medium lacking 2i and LIF but containing 200 nM OHT. At 1 day of differentiation, medium was replaced with medium without OHT. At day 4 of differentiation, the EBs were transferred to gelatinised tissue culture dishes as above.

### KAP1 and RIF1 ChIP

Chromatin immunoprecipitation was performed according to(Bulut-Karslioglu, Perrera et al., 2012). Briefly, for RIF1 and KAP1 ChIP, collected cells were first crosslinked using 2mM disuccinimidyl glutarate (DSG, # BC366 Synchem UG & Co. KG) in PBS for 45 min. at RT while rotating, washed twice in PBS, followed by 10 min. of additional crosslinking in 1% formaldehyde (#252549, Sigma-Aldrich) in cross-linking buffer (50 mM HEPES pH 7.8, 150 mM NaCl, 1 mM EDTA and 500uM EGTA) at RT. Crosslinking was followed by 5 min. quenching in 0.125 M glycine at RT, washed twice in cold PBS and resuspended in lysis buffer (1% SDS, 10 mM EDTA, 50 mM Tris-HCl pH 8.1, supplemented with protease inhibitor cocktail, #11873580 001, Roche). Chromatin fragmentation was performed using Soniprep 150 to produce a distribution of fragments enriched between 300 and 400 bp. The lysate was precleared by centrifugation at low speed 2,000 rpm for 20 min at 4 °C. Chromatin was quantified using Qubit dsDNA High Sensitivity assay kit (#Q32854, Life Technologies). Immunoprecipitation was performed by incubating 100 μg of chromatin diluted in 10 volumes of Dilution buffer (1% Triton X-100, 2 mM EDTA, 167 mM NaCl, 20 mM Tris-HCl pH 8.1, including Protease Inhibitor) overnight rotating at 4 °C together with either α-KAP1 or α-RIF1 antibodies (see Supplementary Table 2) or IgG only control (#sc-2026, Santa Cruz), 10% of chromatin was isolated as input control. The following day, 50 μl of Dynabeads protein G slurry (#10004D, Thermo Fisher) per ChIP sample was added and incubated rotating for another 2 hours at 4 °C. The beads were magnet-separated and washed twice with Low salt buffer (0.1% SDS, 1% Triton X-100, 2 mM EDTA, 150 mM NaCl, 20 mM Tris-HCl pH8.1), one time each with High salt buffer (0.1% SDS, 1% Triton X-100, 2 mM EDTA, 500 mM NaCl, 20 mM Tris-HCl pH8.1), LiCl buffer (0.25M LiCl, 0.5% NP-40, 0.5% sodium deoxycholate, 1mM EDTA,10 mM Tris-HCl pH 8.1) and finally TE. Each wash was performed for 5 min. on a rotating wheel at 4 °C and all buffers were supplemented with protease inhibitor cocktail (#11873580 001, Roche). Prior to elution, samples were rinsed once in TE without protease inhibitor. ChIP-DNA was eluted from the beads by rotating at RT for 1 hour in Elution buffer (1% SDS, 100 mM NaHCO_3_). Beads were separated and the supernatants as well as input samples were subjected to RNAse A (#R5250, Sigma-Aldrich) treatment (1.5µl/sample) for 1 hour at 37 °C followed by decrosslinking using Proteinase K (#P6556, Sigma-Aldrich) treatment (45µg/sample) overnight at 60 °C. The following day, ChIP-DNA and input samples were purified using ChIP DNA Clean and Concentrator kit (#D5205, Zymo Research) and retrieved DNA as well as input DNA was quantified using Qubit dsDNA High Sensitivity assay kit (#Q32854, Life Technologies). The concentration of ChIP-DNA and input samples were adjusted to maintain a similar ratio of ChIP-DNA:INPUT between different ChIP experiments. qPCRs were performed using the SYBR Green reaction mix (#04887352001, Roche) on a LightCycler 96 Instrument (Roche), following standard protocols. Enrichments over input control were calculated for each respective primer set. Primer sequences are presented in Supplementary Table 3.

### RNA extraction, reverse transcription and RT-qPCR

Frozen cell pellets were lysed and homogenised using QIAshredder column (#79656, QIAGEN) followed by RNA extraction using the RNeasy kit (#74106, QIAGEN) according to the manufacturer’s instructions. On column-DNAse treatment was performed at 25 °C-30 °C for 20 min. using RQ1 RNase-Free DNase (#M6101, Promega). After elution, a second round of DNAse treatment was performed using 8U of DNase/sample, incubated at 37 °C for 20 min. The reaction was terminated by adding 1μl of RQ1 DNase Stop Solution and incubated at 65 °C for 10 min. RNA was quantified using Nanodrop, and cDNA synthesis was performed using RevertAid H Minus First Strand cDNA kit (#K1632, Thermo Scientific) using random hexamer priming. qPCRs were performed using the SYBR Green reaction mix (#04887352001, Roche) on a LightCycler 96 Instrument, following standard protocols. Gene expression data was normalised against a geometric mean generated by RT-qPCR of either: *Gapdh, Ubiquitin* and *β-Actin* or *Rplp0, Ubiquitin* and *Sdha*. For flavopiridol or triptolide treated cells, gene expression levels were normalised against *18S* ribosomal RNA. For RT-qPCR analysis of E6.5 embryos derived from *Rif1*^*+/-*^ timed-matings, isolated embryos were individually collected in 200 μL of RLT buffer (QIAGEN RNeasy Kit). 50 μL were used for extraction of genomic DNA (Invitrogen PureLink genomic DNA mini kit). Out of 30 μL eluted genomic DNA, 3 μl were used for genotyping and sex-typing PCRs. The remaining 150 μL of RLT-dissolved embryo were used for RNA extraction (QIAGEN RNeasy Kit), the RNA was quantified and equal amounts were used for reverse transcription. *Gapdh* was used as reference gene. Primers for RT-qPCR are listed in Supplementary Table 4. Absolute expression levels were calculated using the Ct (2^-ΔCt^) method and relative levels using Ct (2^-ΔΔCt^) (Livak & Schmittgen, 2001).

## Supporting information

all suppl

## Acknowledgments

We would like to acknowledge David Kelly from the COIL facility, WTCCB, University of Edinburgh; Emerald Perlas from the Histology Facility of the Epigenetics & Neurobiology Unit, EMBL Rome; Violetta Parimbeni for mouse husbandry, (Epigenetics & Neurobiology Unit, EMBL Rome). We would like to thank Phil Avner (Epigenetics & Neurobiology Unit, EMBL Rome) for advice, reagents, support, discussions and critically reading the manuscript. Rafael Galupa (EMBL Heidelberg) and Jacqueline Mermoud (University of Marburg) are thanked for critically reading the manuscript. Titia de Lange (The Rockefeller University) is thanked for initially supporting the generation of the *Rif1* knockout mice. Joost Gribnau and Cristina Gontan (Erasmus MC, University Medical Center, Rotterdam) are thanked for the Xist-luciferase reporter plasmid. Andrew Jarman (Centre for Discovery Brain Sciences, Edinburgh) and Sally Lowell (MRC Centre for Regenerative Medicine, Edinburgh) are both thanked for providing reagents.

EE received funding from the European Union’s Horizon 2020 research and the Marie Skłodowska-Curie Individual Fellowship grant agreement No. 660985 and from the ERC consolidator award 726130 to SCBB. LP and LB were funded by the ERC consolidator award 726130 to SCBB. RF was funded by the EMBL Interdisciplinary Postdoc (EIPOD) fellowship under Marie Curie Actions (COFUND).

## Author Contributions

EE has created the cellular system, performed the majority of the experiments and co-written the manuscript. RF initiated the project and performed some of the early experiments, like the staining of E3.5 embryos. LP has performed some of the ChIP experiments, the triptolide treatment and the Luciferase assay. LB, AC and NBR have performed KAP1 KD, RNA FISH and the analysis. FC has analysed E6.5 embryos by RT-qPCR and performed the H&E analysis. GK has analysed the RNA seq data, supervised by MV. NBR was supervised by AC, who also critically read the manuscript. AP has isolated and stained the E5.5 embryos. SCBB has conceived the project, performed some of the experiments and written the manuscript.

## EXPANDED VIEW FIGURE LEGENDS

**Figure EV1.**
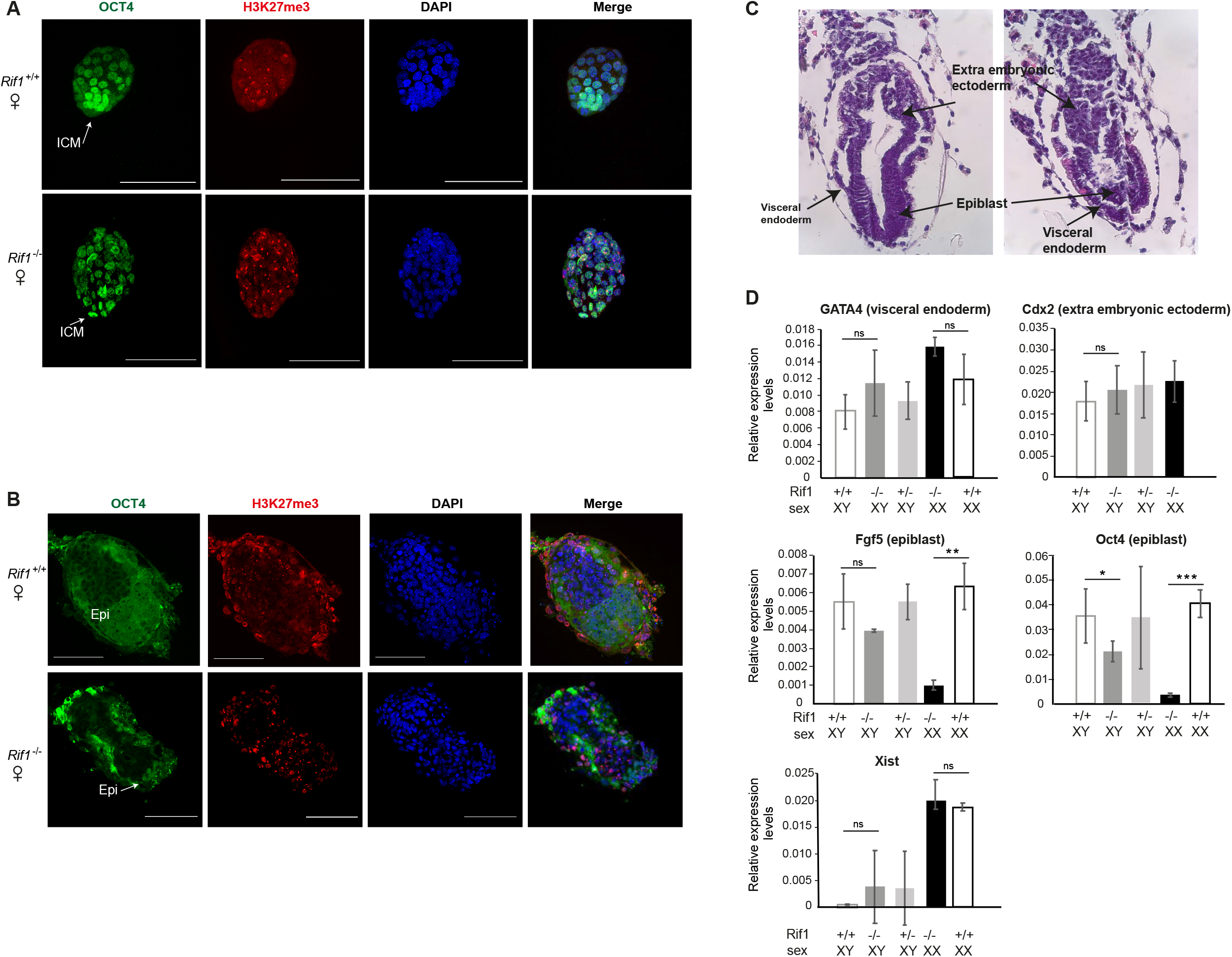
Growth defects in *Rif1*^-/-^ females occur around implantation. Representative whole-mount embryo immunostainings of E3.5 **(A)** and E5.5 **(B)** female embryos. *Rif1*^*+/+*^ *and Rif1*^*-/-*^ embryos are shown. OCT4 (green) marks cells of the inner cell mass (ICM) in **(A)** and of the epiblast (Epi) in **(B)** H3K27me3 (red) labels the Xi. DNA was stained by DAPI (blue). Scale bars =0.1 mm. **(C)**. Sections of E6.5 embryos derived from *Rif1*^*+/-*^ inter-crosses, stained with Hematoxylin and Eosin. On the left an example of a normal-looking embryo, on the right an embryo from the same litter, showing a strongly reduced epiblast. (**D)**. RT-qPCR from RNA extracted from single E6.5 embryos derived from *Rif1*^*+/-*^ inter-crosses. Mean ±standard deviation from a minimum of three biological replicates. *p* was calculated using unpaired Student’s *t* test. *p ≤ 0.05, **p ≤ 0.01, ***p ≤ 0.001 and ns=not significant. The RNA levels of genes expressed in the extra embryonic tissues (*Gata4* for the visceral endoderm and *Cdx2* for the extra embryonic ectoderm) support the conclusion that extra-embryonic tissues are not overtly affected by *Rif1* deficiency. On the contrary, the RNA levels transcribed from genes expressed specifically in the epiblast (*Oct4* and *Fgf5*) are strongly reduced in *Rif1* deficient embryos, more prominently in the females. Xist RNA levels are comparable between *Rif1* wild type and null embryos. Since the epiblast is specifically reduced, the Xist levels detected in *Rif1*^*-/-*^ females indicate that the process of imprinted X inactivation in the extra-embryonic tissues is not affected by *Rif1* deficiency, a conclusion supported by the presence of H3K27me3 in the extra embryonic tissues in **(B)**.

**Figure EV2.**
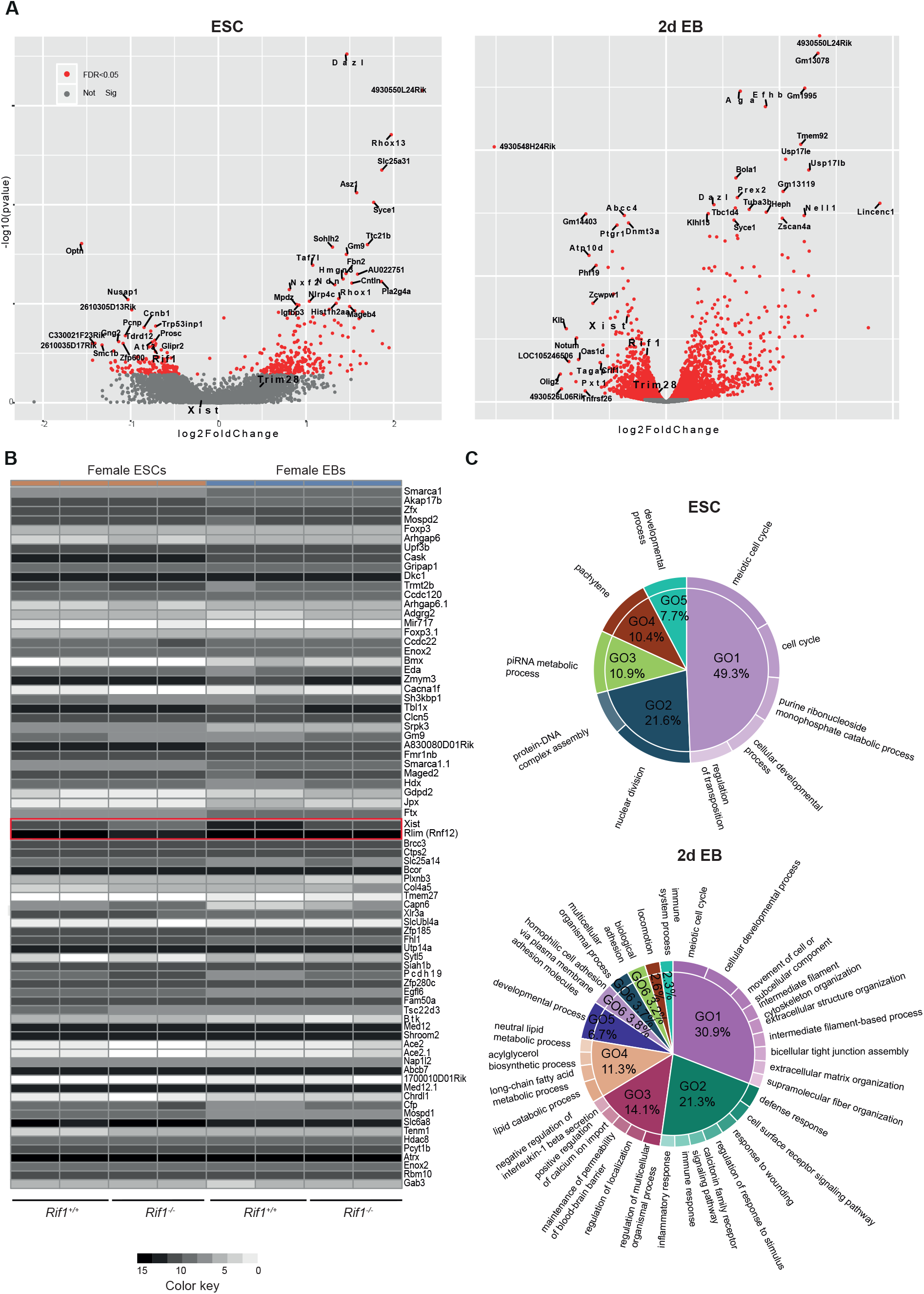
Transcriptome analysis in *Rif1* null ESCs and early-stage EBs. **(A)**. Volcano plot summarising the top differentially expressed genes in *Rif1*^*-/-*^ (*Rif1*^*F/F*^ +OHT) ESCs (left) and EBs (right), compared to *Rif1*^*+/+*^ *(Rif1*^+/+^+OHT). Two independent cell lines were analysed per genotype. In red, genes whose differential expression is statistically significative (FDR<0.005). *Trim28* (*Kap1*) expression is not significantly changed in *Rif1* null cells, whilst *Xist* expression levels are significantly down regulated in EBs. **(B)**. Heat-maps summarising the logarithmically transformed, normalised expression levels of randomly-chosen X-linked genes in *Rif1*^*+/+*^ and *Rif1*^*-/-*^ ESCs (brown) and 2 days EBs (blue). Failure to upregulate *Xist* in *Rif1*^*-/-*^ EBs is specific, as other X-linked genes do not show the same behaviour. *Rlim* (*Rnf12*) expression, for example, is not affected by *Rif1* deletion (highlighted, along with Xist, in the red box). **(C)**. Pie charts summarising the biological processes (GO enrichment analysis, GOrilla) most represented among the genes whose expression is significantly deregulated upon *Rif1* deletion in ESCs (top) and EBs (bottom). Differentially expressed genes were obtained from the DESeq2 (adjusted p-value, 0.05 and llog2FCl>0.5). Genes linked to developmental processes (GO5) represent only a small percentage of differentially expresses genes (about 50% upregulated and 50% downregulated).

**Figure EV3.**
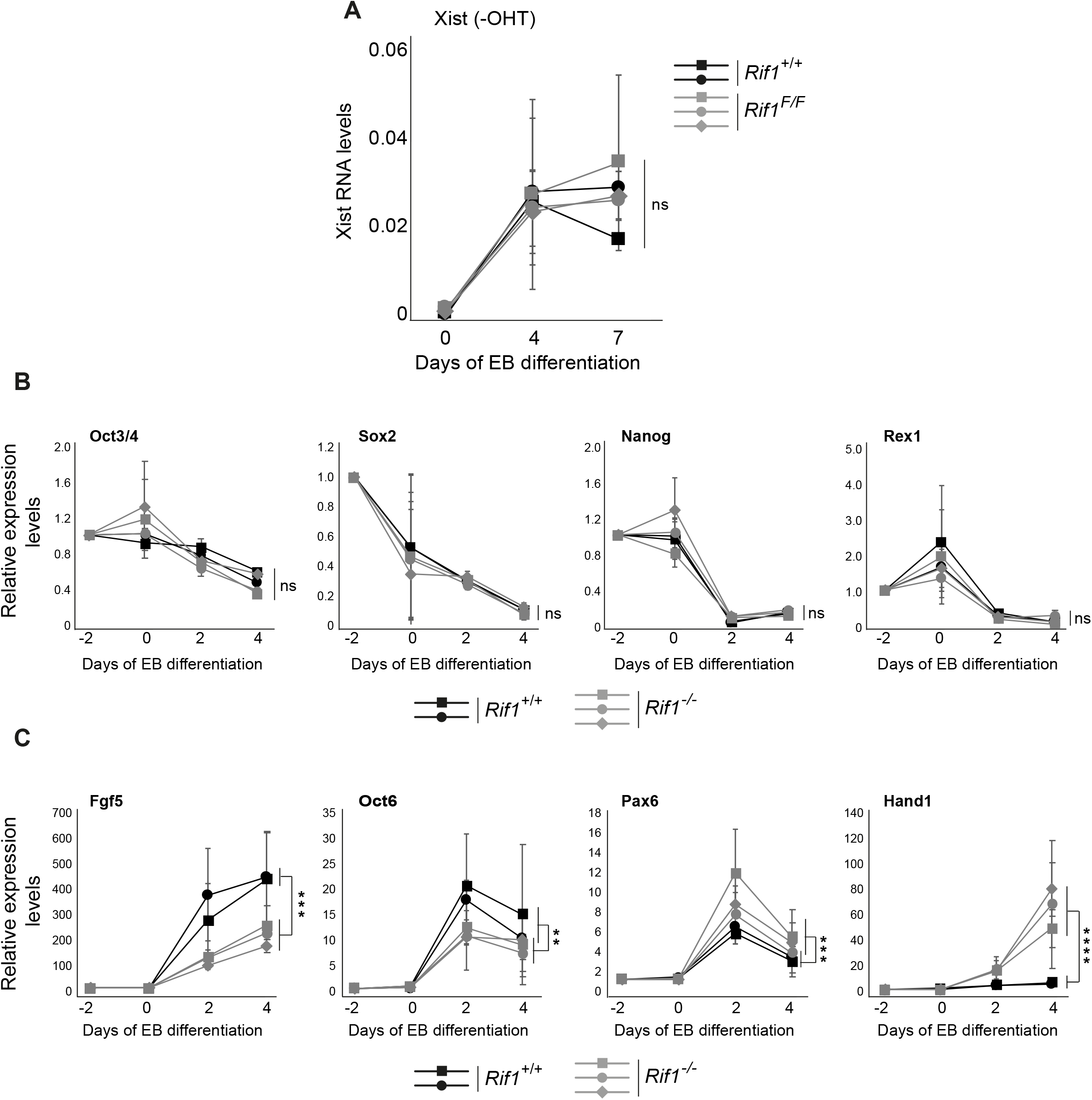
*Rif1*^*-/-*^ mESCs can exit pluripotency and commit to differentiation. **(A)**. Xist RNA levels monitored by RT-qPCR in two independent *Rif1*^*+/+*^ (black) and three independent *Rif1*^*F/F*^ (grey) female mESC lines, during EB differentiation, in the absence of OHT. Xist RT-primers Xist ex7 F and R were used. Values are normalised to a geometric mean consisting of the expression of *Gapdh, Ubiquitin* and *β-Actin*. Expression levels are plotted as mean ±standard deviation of a minimum of three individual experiments with statistical significance determined using 2-way ANOVA. ns=not significant. **(B and C)**. Time course analysis of expression levels for the indicated genes, during EB differentiation of two individual *Rif1*^*+/+*^ (*Rif1*^*+/+*^ +OHT, black) and three individual *Rif1*^*-/-*^ (*Rif1*^*F/F*^ +OHT, grey) female mESC lines. Expression levels are first normalised to a geometric mean consisting of the expression of *Gapdh, Ubiquitin* and *β-Actin* and then plotted relative to pre-samples (day-2), as mean ±standard deviation of three or four individual experiments with statistical significance determined using 2-way ANOVA (**p ≤ 0.01, ***p ≤ 0.001, ****p ≤ 0.0001 and ns=not significant) comparing *Rif1*^*+/+*^ to *Rif1*^*-/-*^ cells. **(B)**. Pluripotency-associated genes and **(C)** genes expressed during early-stages of differentiation.

**Figure EV4.**
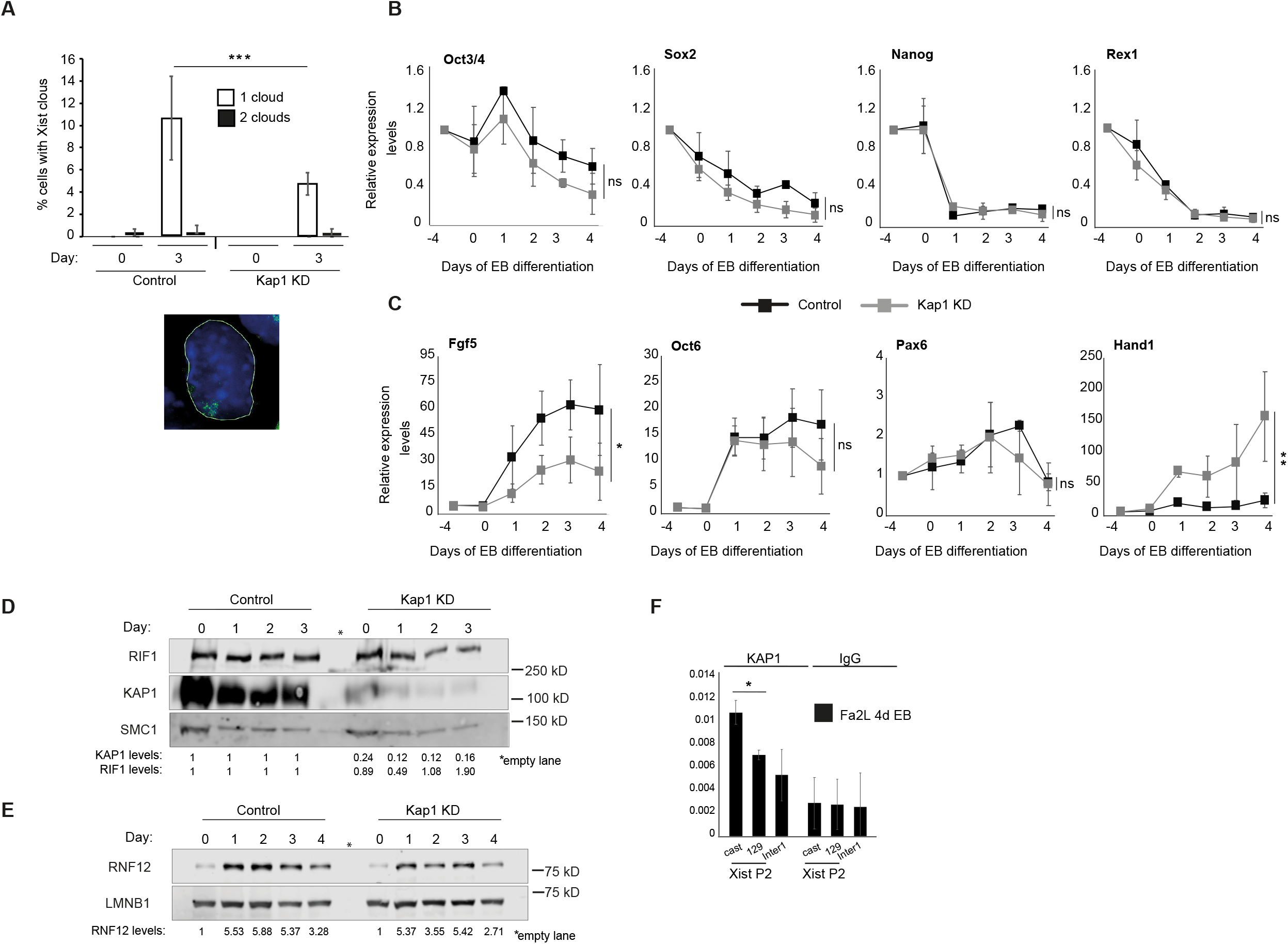
Exit from pluripotency, differentiation, RNF12 upregulation and *Tsix* downregulation in KAP1 knock down cells. **(A)**. Quantification of Xist cloud formation (1 or 2, open or filled bars, respectively) by RNA FISH in Control and Kap1 KD female mESCs, differentiated into EBs for 3 days. The average of two experiments is shown. N>400 cells. The error bars represent ± standard deviation. *p* was calculated by 𝒳^2^ (***p ≤ 0.001). Below, an example of Xist cloud RNA FISH signal. Green=Xist, blue=DAPI. **(B and C)**. Time course analysis of gene expression levels quantified by RT-qPCR during EB differentiation of female mESCs infected with shRNA directed against Luciferase (control-black) and KAP1 (grey), at the indicated timepoints. Values have first been normalised to a geometric mean consisting of the expression *Rplp0, Ubiquitin* and *Sdha* and then presented relative to pre-samples (day: -4) Mean ±standard deviation from three independent experiments are shown. Statistical significance was determined using 2-way ANOVA (*p ≤ 0.05, **p ≤ 0.01 and ns=not significant). **(B)**. pluripotency-associated genes and genes expressed during early-stages of differentiation. **(D)**. Representative western blot analysis of RIF1 and KAP1 levels in Kap1 KD female mESCs. SMC1: loading control. Quantification of RIF1 and KAP1 protein levels normalised to SMC1 and relative to control cells are shown below. **(E)**. Representative western blot analysis of RNF12 levels in proteins extracted from female mESC infected with shRNA directed against Luciferase (control) and KAP1, at the indicated timepoints during EB differentiation. LMNB1: loading control. Below, quantification of RNF12 protein levels relative to day 0 of control and KAP1 KD cells respectively. Values normalised to LMNB1. **(F)**. Using allele-specific primers, ChIP-qPCR was used to analyse the association of KAP1 with *Xist* P2 in differentiating Fa2L cells. *cast* indicates association with the *castaneus Xist* P2 and *129* indicates association with the *129 Xist* P2. Enrichments are presented relative to input DNA. Mean ±standard deviation from a minimum of three independent experiments. *p* was calculated by Student’s two-tailed paired *t* test (*p ≤ 0.05).

**Figure EV5.**
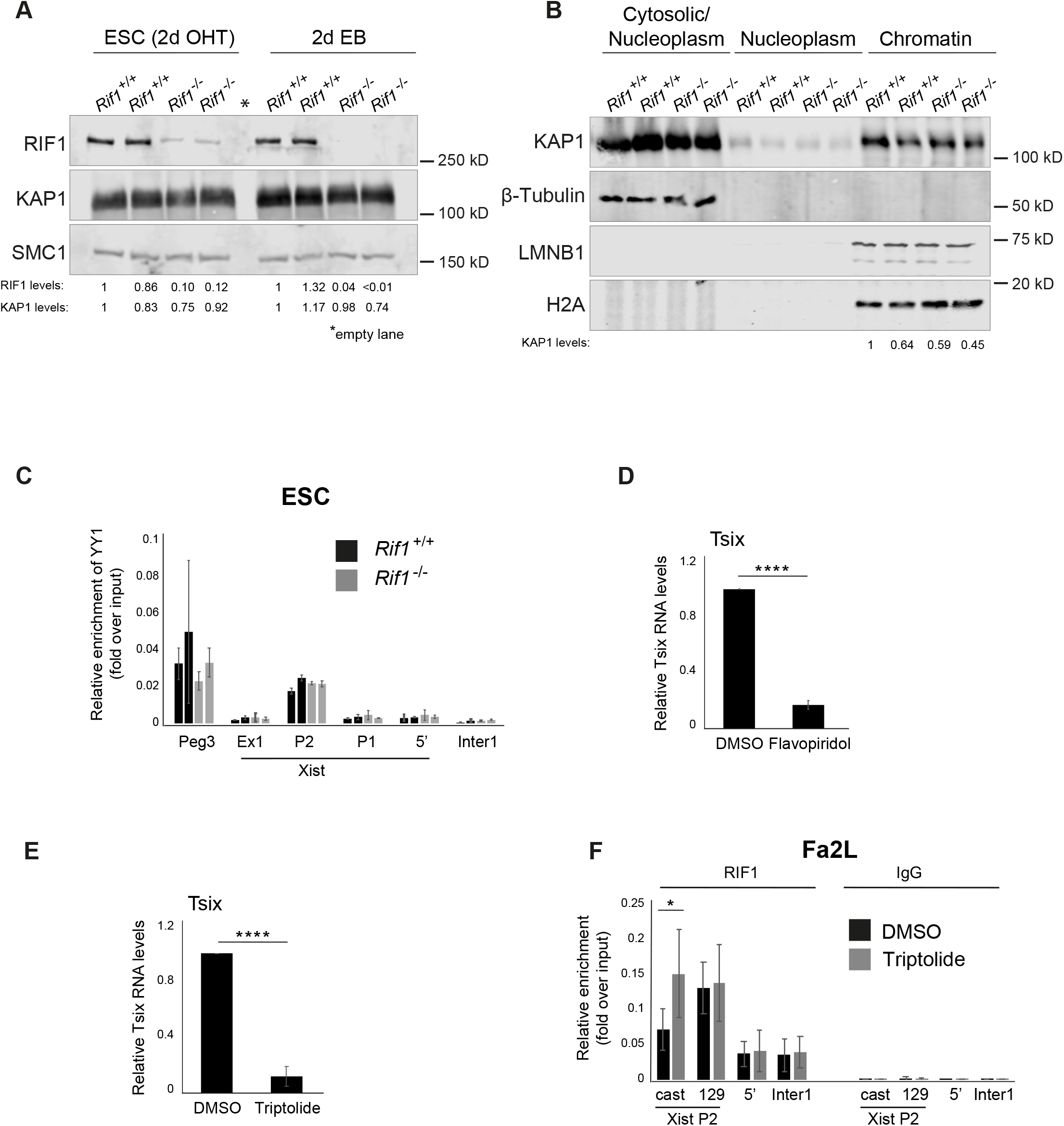
The overall binding of KAP1 to chromatin is unaffected in *Rif1*^*-/-*^ mESCs. **(A)**. RIF1 and KAP1 levels analysed by western blot in protein extracts from two *Rif1*^*+/+*^ (*Rif1*^*+/+*^ +OHT) and two *Rif1*^*-/-*^ (*Rif1*^*F/F*^ +OHT) independent female mESC lines, following 2 days of OHT treatment and at 2 days EB. SMC1: loading control. Below, quantifications of RIF1 and KAP1 protein levels shown as relative levels compared to one of the *Rif1*^*+/+*^ cells. Values normalised to SMC1. **(B)**. Western blot analysis of KAP1 levels in protein extracts from the indicated cell fractions, from two *Rif1*^*+/+*^ (*Rif1*^*+/+*^ +OHT) *and* two *Rif1*^*-/-*^ (*Rif1*^*F/F*^ +OHT) independent female mESC lines, following 2 days of OHT treatment. β-tubulin: marker for the cytosolic fraction. LNMB1 and histone H2A: markers for the chromatin/insoluble fraction. Below, quantifications of KAP1 protein levels detected in the chromatin fractions shown as relative levels compared to one of the control cells. Values normalised to H2A. **(C)**. YY1 association with the *Xist* promoter in two independent *Rif1*^*+/+*^ (*Rif1*^*+/+*^ +OHT, black) and two *Rif1*^*-/-*^ (*Rif1*^*F/F*^+OHT, grey) female mESC lines, analysed by ChIP-qPCR. Ex1 indicates a region within Xist exon 1, 2.5 kb downstream of *Xist* transcriptional start site (TSS), P2 indicates *Xist* promoter P2 spanning the YY1 consensus motif. Peg3 indicates a known YY1-associated site on the *Peg3* gene. Tsix RNA levels following flavopiridol and triptolide **(E)** treatment, relative to DMSO treated Fa2L cells. RNA levels were first normalised to *18S* ribosomal RNA and plotted as mean from three individual experiments ±standard deviations. Statistical significance was determined using Student’s two-tailed unpaired *t* test (****p ≤ 0.0001). **(F)**. Allele-specific RIF1 association with *Xist* P2 in Fa2L cells following treatment with DMSO only (black) or triptolide (grey). *cast* indicates association with the *castaneus Xist* P2 promoter and *129* indicates association with the *129 Xist* P2 promoter. Enrichments are presented relative to input DNA. Mean ±standard deviation of three independent experiments. *P* calculated by Student’s two-tailed paired *t* test. (*p ≤ 0.05 and ns=not significant).

**FIGURE S1.**
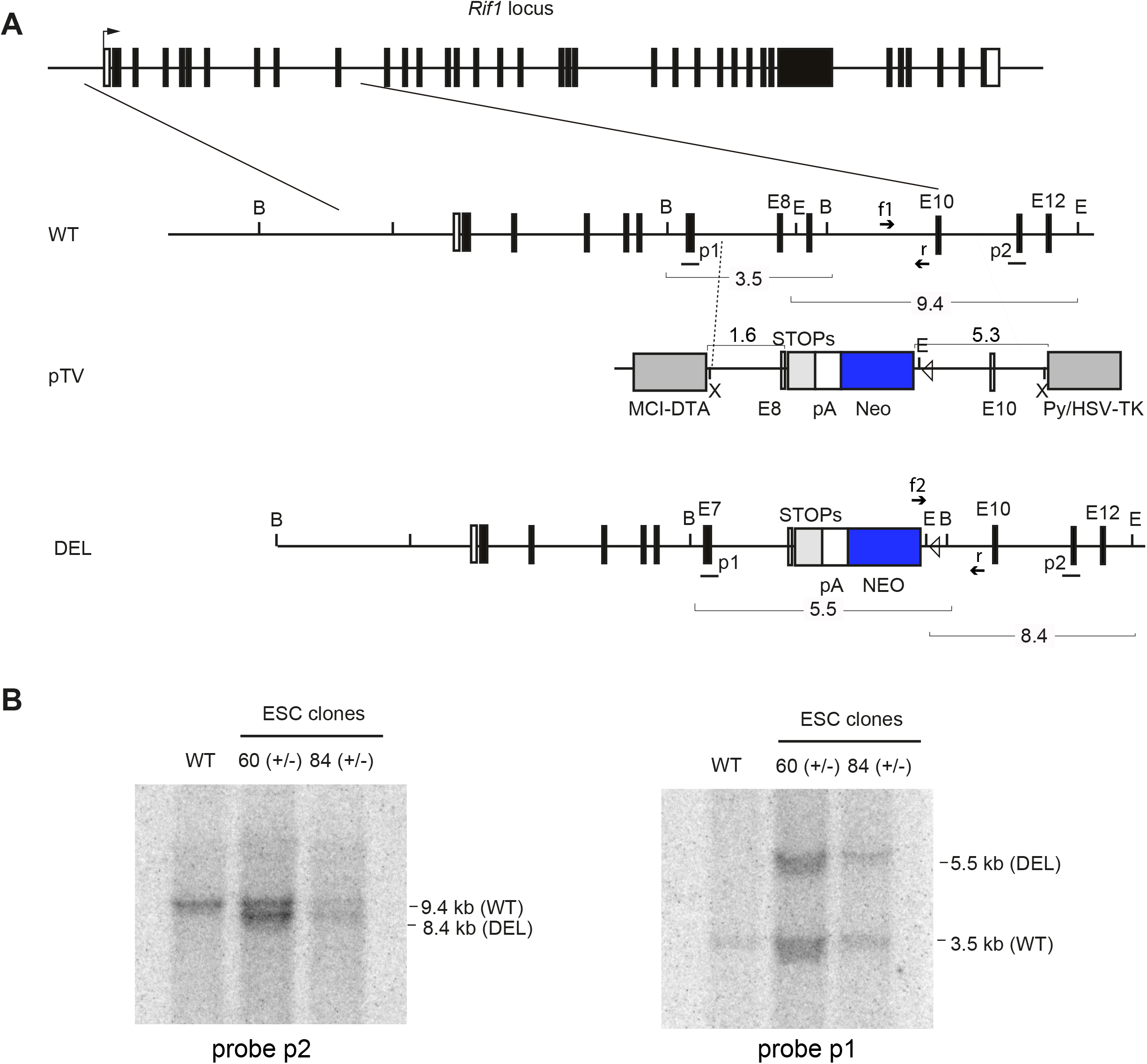

**FIGURE S2.**
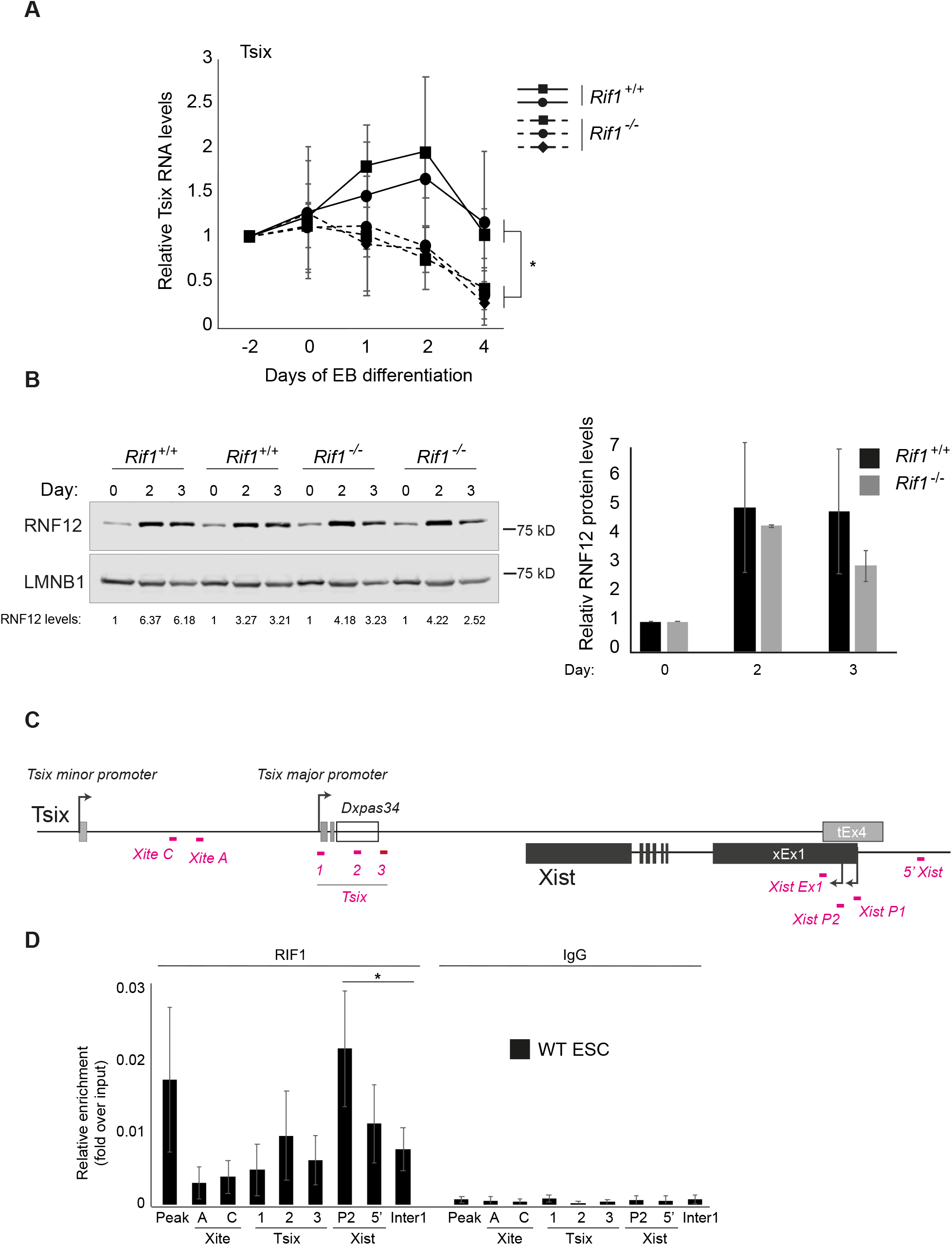

**FIGURE S3.**
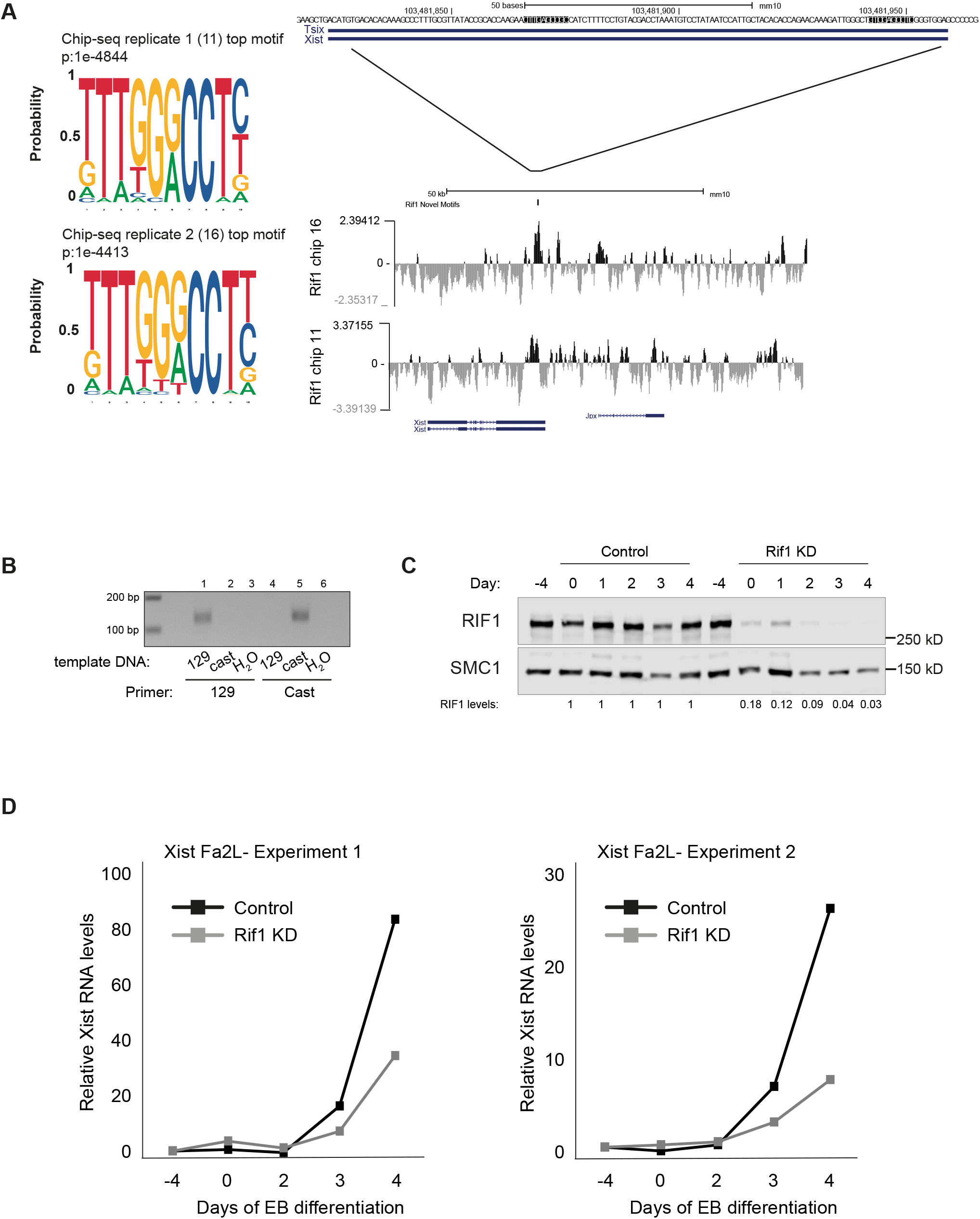

**FIGURE S4.**
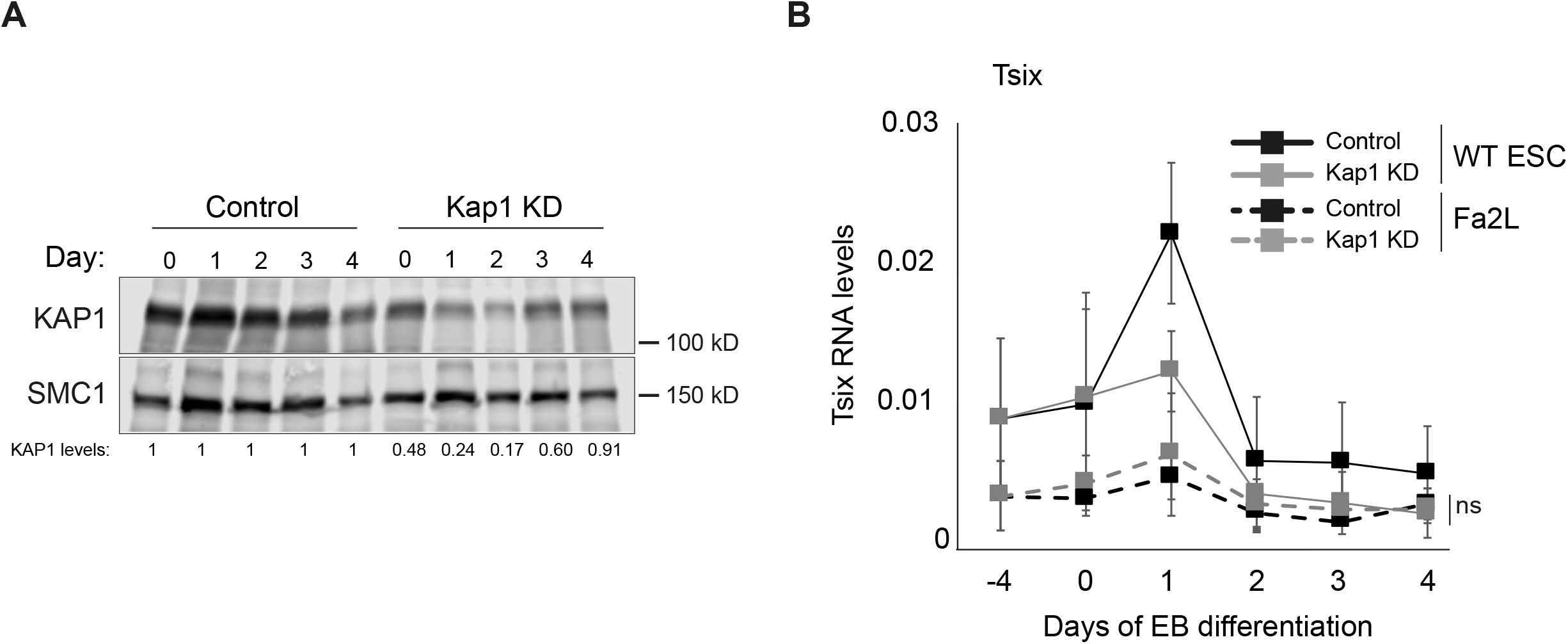

