## Supplementary material for "A RIF1/KAP1-based toggle switch stabilises the identities of the inactive and active X chromosomes during X inactivation": all suppl

**Supplementary information:**

Supplementary figure legends

Supplementary Figures

Supplementary Tables

Supplementary methods

**Figure S1. RIF1 targeting strategy.**

**(A).** Schematic diagram of the mouse *Rif1* locus (WT), the targeting vector (pTV), the *Rif1* null allele (DEL), consisting of a STOP-pA-NEO (grey, white and blue rectangles), replacing exons 8 and 9. A lox P site flanking the NEO cassette is indicated by a triangle. Fragment sizes are indicated for each genotype, and the probes p1 and p2 are shown. f1, f2 and r, primers for genomic PCR; B, BsrGI; E, EcoRV; X, XhoI.

**(B).** Southern blots of targeted mESC clones injected into blastocysts to obtain chimeric mice. Genomic DNA digested with BsrGI and probed with p1 (right) or with EcoRV and probed with p2 (left).

**Figure S2. The failed *Xist* upregulation at the onset of differentiation in *Rif1*$^{-/-}$ mESCs is not caused by lack of RNF12 upregulation or defective *Tsix* downregulation.**

**(A).** Tsix RNA levels in two independent *Rif1*$^{+/+}$ female lines (*Rif1*$^{+/+}$ +OHT, solid line) and three independent *Rif1*$^{-/-}$ female lines (*Rif1*$^{F/F}$ +OHT, dashed line) at indicated timepoints, during EB differentiation. Expression levels are first normalised to a geometric mean consisting of the expression of *Gapdh, Ubiquitin* and *β-Actin* and then plotted relative to pre-sample levels (day -2), as mean ±standard deviation of three individual experiments, with the exception of one of the *Rif1*$^{-/-}$ lines, which was only included in two of the experiments. Statistical significance was determined using 2-way ANOVA, comparing *Rif1*$^{+/+}$ to *Rif1*$^{-/-}$ (*p ≤ 0.05). **(B).** RNF12 levels analysed by western blot of protein samples from two independent *Rif1*$^{+/+}$ (*Rif1*$^{+/+}$ +OHT), and two independent *Rif1*$^{-/-}$ (*Rif1*$^{F/F}$ +OHT) female mESC lines, at indicated timepoints, during EB differentiation. LAMIN B1 (LMNB1): loading control. Right, quantification of RNF12 protein levels relative to day 0. Bar plot shows the mean ±standard deviation of the relative RNF12 levels of the two biological replicates run in the western blot shown. Values normalised to LMNB1. **(C).** Illustration of the Tsix/*Xist* locus with Chip-qPCR primer positions indicated (magenta). **(D).** RIF1 association with different regions known to be important for XCI, assessed by ChIP-qPCR in a WT female mESC line. Peak indicate a previously identified RIF1 positive region. Xite A and C indicate two regions within the Tsix enhancer *Xite*, Tsix region 1 indicates Tsix major promoter, Tsix region 2 indicates the Dxpas34 region, Tsix region 3 indicate a region slightly downstream of the Dxpas34 region. P1 and P2 indicate the two *Xist* promoters, 5' indicates a region 2 kb upstream of *Xist* TSS and inter1 is an intergenic region.Mean ±standard deviation from a minimum of three independent experiments are displayed. *p* calculated by Student's two-tailed paired *t* test comparing RIF1 association to Xist P2 versus intergenic region. ns= not significant.

**Figure S3. *Xist* upregulation is impaired in differentiating Fa2L cells upon RIF1 knock down.**

**(A).** Identification of consensus sequence motifs within RIF1 enriched regions. The motifs were identified by HOMER, using data from two RIF1 ChIP-seq (called peaks, replicate 1 and 2), performed on independently derived ESCs (Foti et al., 2016). The RIF1 peaks within the *Xist* region are shown. In the zoom-in, highlighted in black, the positions of the motif, identified twice within the *Xist* P2 sequence. **(B).** Allele-specific qPCR amplifying *Xist* P2. *129* and *cast* indicate SNP-specific primer pairs. Equal amount of template DNA and primers were used in all reactions. **(C).** Western blot analysis of RIF1 levels in protein samples collected from Fa2L cells infected with Luciferase (Control) and RIF1 shRNA respectively, at the indicated timepoints, following EB differentiation. SMC1: loading control. Below, quantifications of RIF1 protein levels shown as relative levels compared to control cells at the corresponding time points. Values normalised to SMC1. **(D).** *Xist* upregulation relative to RNA levels in the pre-infection sample (day -4), during EB differentiation in Fa2L cells infected with shRNA directed against Luciferase (control-black) and RIF1 (grey), at indicated timepoints. Two individual experiments are shown, left and right. *Xist* primers Xist ex3 F and Xist ex4 R were used. For both experiments, values are normalised to a geometric mean consisting of the expression of *Rplp0, Ubiquitin* and *Sdha.*

**Figure S4. KAP1 is dispensable for *Xist* expression in the Fa2L cells.**

**(A).** Representative western blot analysis of KAP1 levels in protein extracts from Fa2L cells infected with shRNA directed against Luciferase (control) and KAP1 at indicated timepoints during EB differentiation. SMC1: loading control. Below, quantifications of KAP1 protein levels shown as relative levels compared to control cells at the corresponding time points. Values normalised to SMC1. **(B).** RT-qPCR quantification of Tsix RNA levels in a control female mESC line (solid line) and in the Fa2L cell line (dashed line) following infection with shRNA directed against Luciferase (control-black) and KAP1 (grey), at indicated timepoints during EB differentiation. The data from the control mESC line are the same as presented in Fig. 4D. The experiments in the two cell lines presented were conducted in parallel and are presented together here to allow the evaluation of the relative amounts and the dynamics of Tsix upregulation in both cell lines. Expression levels were first normalised to a geometric mean consisting of the expression *Rplp0, Ubiquitin* and *Sdha* and presented as mean ±standard deviation from a minimum of three independent experiments. Statistical significance was determined using 2-way ANOVA comparing Tsix levels in control treated Fa2L cells versus Kap1 knock down Fa2L cells. ns=not significant.

**FIGURE S1**

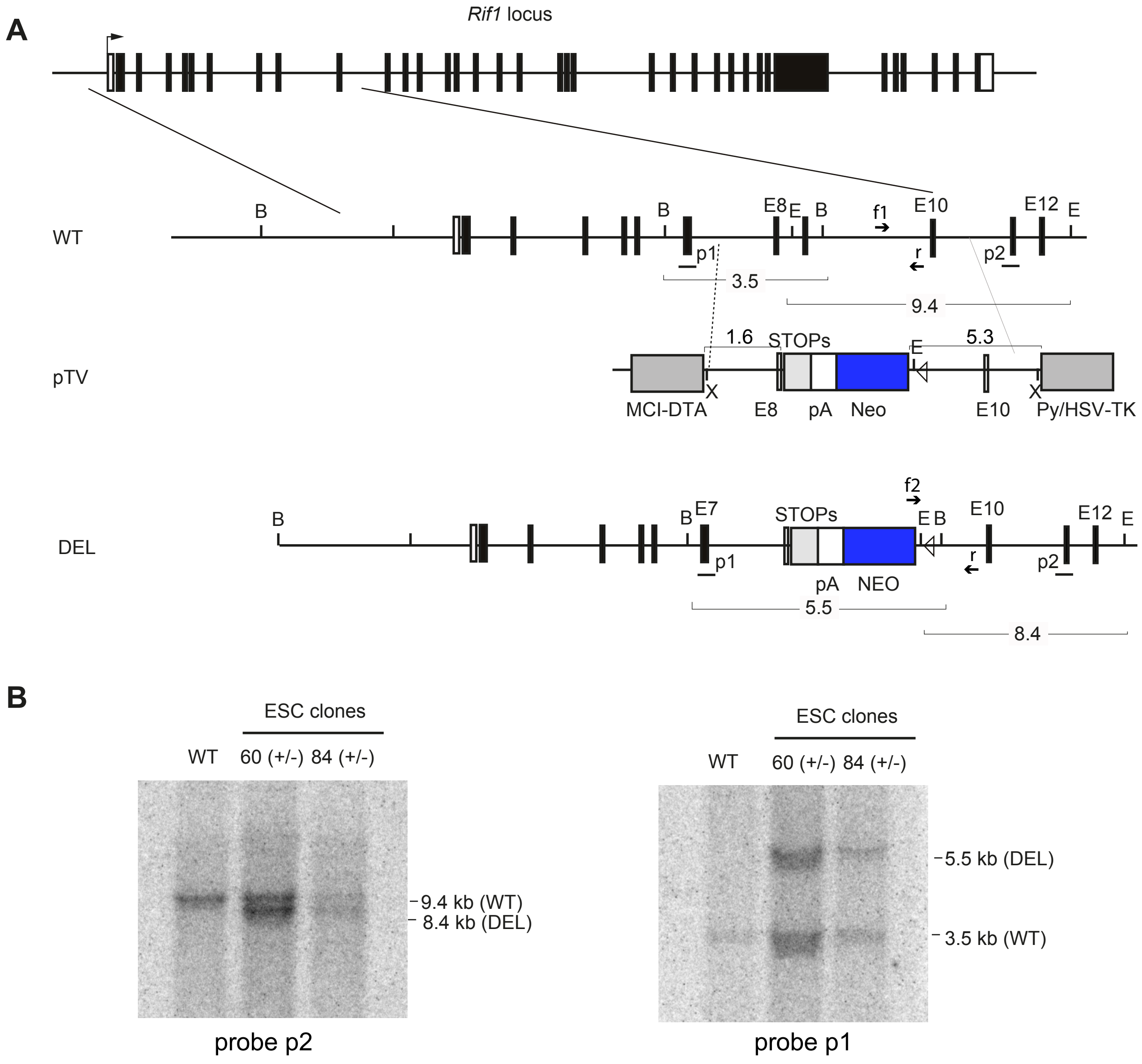
A
*Rif1* locus
WT
B
B
E8
E
B
f1
E10
E12
E
p1
r
p2
3.5
9.4
pTV
1.6
STOPs
E
5.3
X
X
MCI-DTA
E8
pA
Neo
E10
Py/HSV-TK
DEL
f2
B
B
E7
STOPs
E
B
E10
E12
E
p1
r
p2
pA
NEO
5.5
8.4
B
ESC clones
WT
60 (+/-)
84 (+/-)
9.4 kb (WT)
8.4 kb (DEL)
probe p2
ESC clones
WT
60 (+/-)
84 (+/-)
5.5 kb (DEL)
3.5 kb (WT)
probe p1

#### FIGURE S2

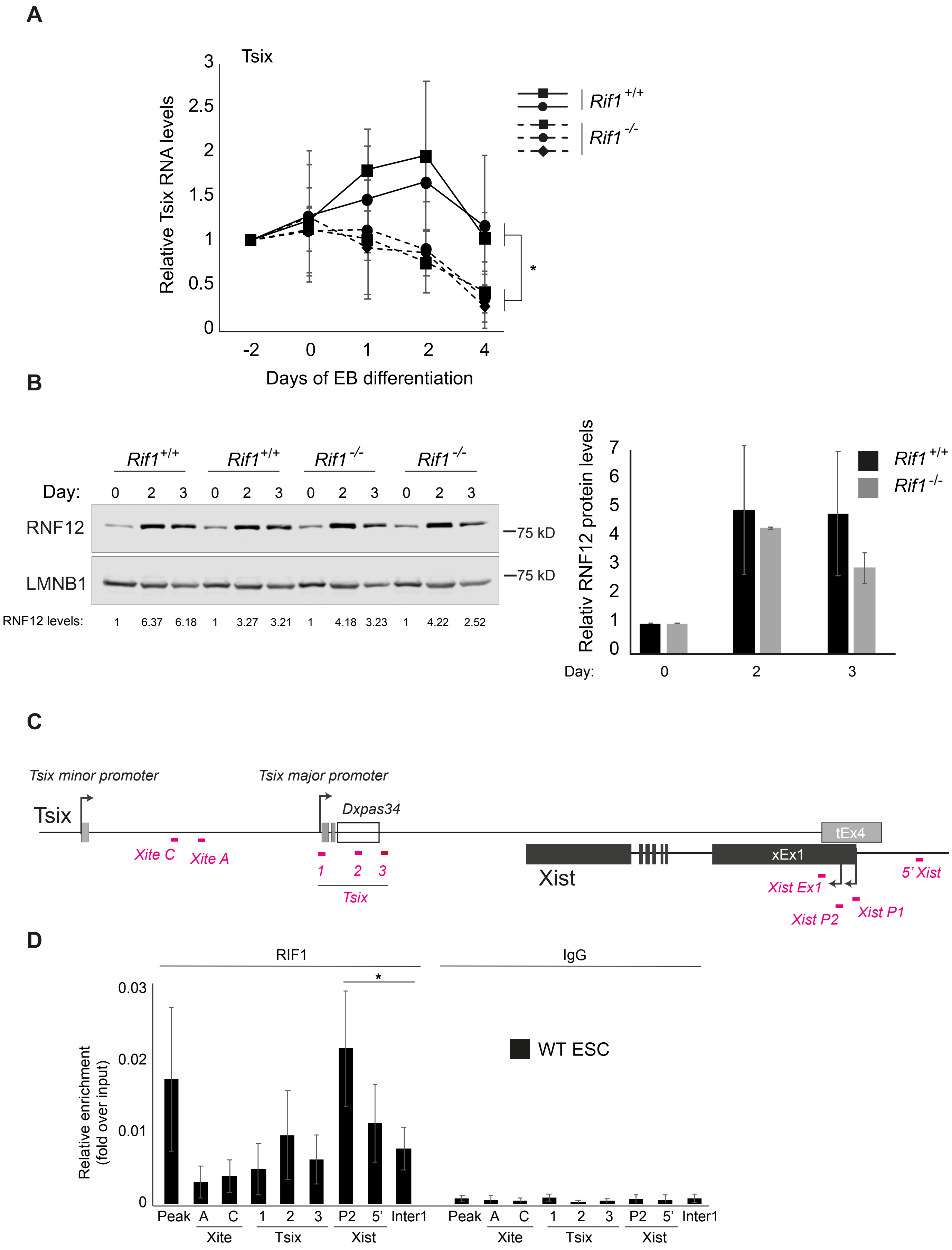

A
Tsix
Relative Tsix RNA levels
3
2.5
2
1.5
1
0.5
0
-2
0
1
2
4
Days of EB differentiation
Rif1 +/+
Rif1 -/-
*
B
Rif1 +/+
Rif1 +/+
Rif1 -/-
Rif1 -/-
Day: 0 2 3 0 2 3 0 2 3 0 2 3
RNF12
LMNB1
—75 kD
—75 kD
RNF12 levels: 1 6.37 6.18 1 3.27 3.21 1 4.18 3.23 1 4.22 2.52
Relativ RNF12 protein levels
7
6
5
4
3
2
1
0
Day: 0 2 3
Rif1 +/+
Rif1 -/-
C
Tsix minor promoter
Tsix major promoter
Tsix
Dxpas34
tEx4
Xite C
Xite A
1 2 3
Tsix
Xist
xEx1
Xist Ex1
5' Xist
Xist P2
Xist P1
D
RIF1
IgG
*
Relative enrichment (fold over input)
0.03
0.02
0.01
0
WT ESC
Peak A C 1 2 3 P2 5' Inter1
Xite Tsix Xist
Peak A C 1 2 3 P2 5' Inter1
Xite Tsix Xist

### FIGURE S3

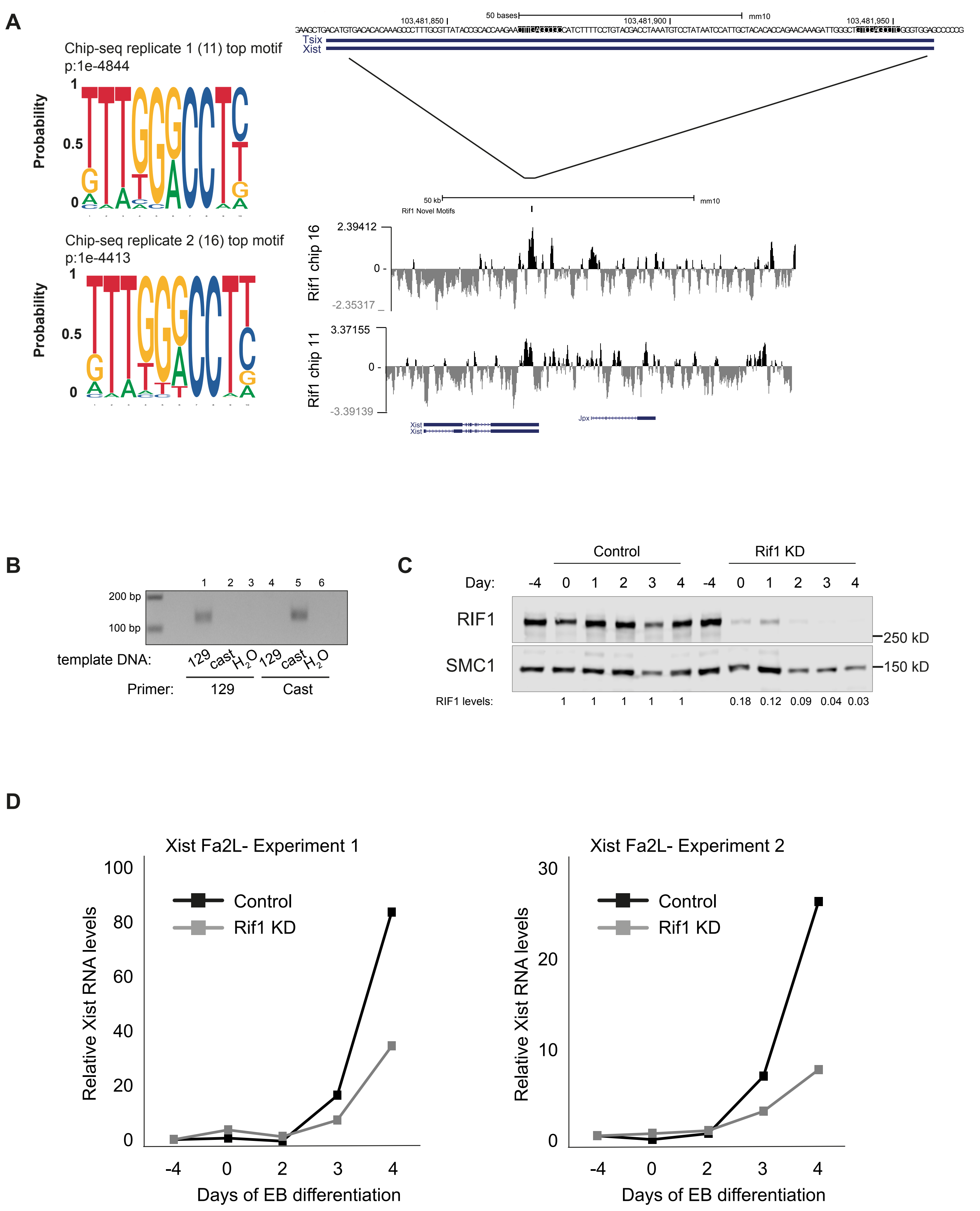
A
Chip-seq replicate 1 (11) top motif
p:1e-4844
Probability
Chip-seq replicate 2 (16) top motif
p:1e-4413
50 bases
mm10
Tsix
Xist
50 kb
Rif1 Novel Motifs
2.39412
-2.35317
Rif1 chip 16
3.37155
-3.39139
Rif1 chip 11
Jpx
B
200 bp
100 bp
template DNA:
129 cast H2O 129 cast H2O
Primer:
129
Cast
C
Control
Rif1 KD
Day:
RIF1
SMC1
250 kD
150 kD
RIF1 levels:
1 1 1 1 1 0.18 0.12 0.09 0.04 0.03
D
Xist Fa2L- Experiment 1
Xist Fa2L- Experiment 2
Control
Rif1 KD
Relative Xist RNA levels
Days of EB differentiation

**FIGURE S4**

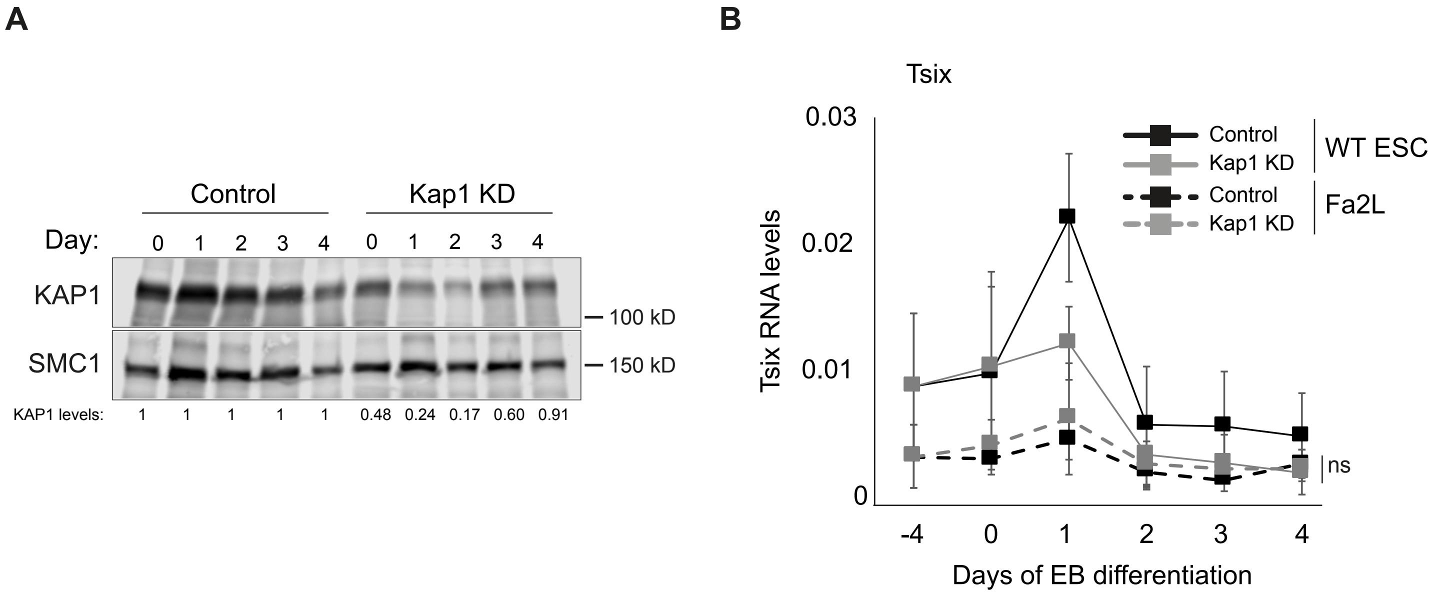
A
Control
Kap1 KD
Day:
0 1 2 3 4 0 1 2 3 4
KAP1
— 100 kD
SMC1
— 150 kD
KAP1 levels:
1 1 1 1 1 0.48 0.24 0.17 0.60 0.91
B
Tsix
Control
Kap1 KD
WT ESC
Control
Kap1 KD
Fa2L
Tsix RNA levels
0.03
0.02
0.01
0
-4 0 1 2 3 4
Days of EB differentiation
ns

### Supplementary Tables

#### Supplemental Table 1

shRNA sequences

| **shRNA** | **Sequence 5' – 3'** | **Description** | **Target** | **Catalogue number/ Reference** |
|---|---|---|---|---|
| LUC F | CCGGATGTTTACTACACTCGGATATCTCGA GATATCCGAGTGTAGTAAACATTTTTT | Control | Luciferase | #SHC007 Sigma-Aldrich |
| LUC R | AATTAAAAAATGTTTACTACACTCGGATA TCTCGAGATATCCGAGTGTAGTAAACA | Control | Luciferase | |
| mRif1-39 F | CCGGCCCTCTATGATCCGAGAAATACTCG AGTATTTCTCGGATCATAGAGGGTTTTTT | Rif1 KD | mouse Rif1 | TRCN0000071339 (The Broad Institute) |
| mRif1-39 R | AATTAAAAAACCCTCTATGATCCGAGAAA TACTCGAGTATTTCTCGGATCATAGAGGG | Rif1 KD | mouse Rif1 | |
| mRif1-40 F | CCGGGCACCTTATGACTACTAAATTCTC GAGAATTTAGTAGTCATAAGGTGCTTTTTT | Rif1 KD | mouse Rif1 | TRCN0000071340 (The Broad Institute) |
| mRif1-40 R | AATTAAAAAAGCACCTTATGACTACTAA ATTCTCGAGAATTTAGTAGTCATAAGGTGC | Rif1 KD | mouse Rif1 | |
| mKap1-F | CCGGCCGCATGTTCAAACAGTTCAACTCG AGTTGAACTGTTTGAACATGCGGTTTTT | Kap1 KD | mouse Kap1 | (Ding *et al*, 2018) |
| mKap1-R | AATTAAAAACCGCATGTTCAAACAGTTCAA CTCGAGTTGAACTGTTTGAACATGCGG | Kap1 KD | mouse Kap1 | |

#### Supplemental Table 2

Antibodies

| **Primary Antibody** | **Catalogue Number** | **Company/Ref** | **Application/Concentration** | | |
|---|---|---|---|---|---|
| | | | **Western Blot** | **IF** | **ChIP** |
| **RIF1** Polyclonal rabbit | #1240 | (Buonomo *et al*, 2009) | 1:2000- 1:5000 | N/A | 1.5µl/100µg chromatin |
| **SMC1** Polyclonal rabbit | A300-055A | Bethyl Laboratories | 1:3000 | N/A | N/A |
| **RNF12** Polyclonal mouse | H00051132-B01P | Abnova | 1:1000 | N/A | N/A |
| **KAP1** Polyclonal rabbit | ab10484 | Abcam | 1:3000 | N/A | N/A |
| **KAP1 20C1** Monoclonal mouse | ab22553 | Abcam | N/A | N/A | 15µg/100µg chromatin |
| **YY1** Rabbit polyclonal | 61779 | Active Motif | N/A | N/A | 10µg/100µg chromatin |
| **LaminB1** Polyclonal rabbit | ab16048 | Abcam | 1:1000 | N/A | N/A |
| **β-Tubulin** Monoclonal mouse | T4026 | Sigma | 1:1000 | N/A | N/A |
| **H2A** Rabbit polyclonal | ab18255 | Abcam | 1:1000 | N/A | N/A |
| **H3** Rabbit polyclonal | 4499 | Cell Signaling | 1:5000 | N/A | N/A |
| **H3K27me3** Rabbit polyclonal | 07-449 | Millipore | N/A | 1:800 | N/A |

#### Supplemental Table 3

ChIP-qPCR primers

| **Primer** | **Sequence 5' – 3'** | **Name in figure(s)** | **Description** | **Tm (°C)** | **Reference** |
|---|---|---|---|---|---|
| Xist P2 F | TCATGTGACCTGCCCTCTAGT | P2 (Xist) | Xist P2 promoter | 60 | (Navarro *et al*, 2005) |
| Xist P2 R | CACCCTACCATAATGCACCA | | | | |
| Xist P1 F | AAGGCTTGGTGGTAGGGGA | P1 (Xist) | Xist P1 promoter | 60 | (Elsasser *et al*, 2016) |
| Xist P1 R | TTTGCTCGTTTCCCGTGGAT | | | | |
| Xist_P2_129 F | AGCGGACTGGATAAAAGCAA**C** | 129 | Xist P2 promoter *129*-specific | 67 | This paper |

| | | | | | |
|---|---|---|---|---|---|
| Xist_P2_cast F | AGCGGACTGGATAAAAGCAA**T** | cast | Xist P2 promoter *castaneus*-specific | 67 | This paper |
| Xist_P2_129/cast R | TACTTCCGGCTAGCACAACC | | Xist P2 promoter | 67 | This paper |
| Xist_P2_YY1 F | TCGACAGCCCAATCTTTGTTC | P2 (Figure S9C) | Spanning YY1 consensus motif | 60 | This paper |
| Xist_P2_YY1 R | ACTTGAGCCGCCATCTTTTC | | | | |
| 5'Xist F | ACCATCAGGTTCGTCAGAGC | 5' | 2 kb upstream of Xist TSS | 60 | This paper |
| 5'Xist R | AAGCAGCTCGTGGGTAGAAC | | | | |
| Xist Ex1 F | AGCAACAAAAGCAAAGCCTG | Ex1 | 2.5 kb downstream of Xist TSS | 60 | This paper |
| Xist Ex1 R | TCATCCACCGAGCTACTCTTC | | | | |
| Jpx F | TTAAAGACGGGCAAGAAACG | Jpx | Jpx promoter | 60 | This paper |
| Jpx R | CTGTCCCTAAATGGCTGCTC | | | | |
| Ftx TSS1 F | TAATGCGCAACATCCTTTTG | TSS1 (Ftx) | | 60 | This paper |
| Ftx TSS1 R | CAGCAGGAGGTAAAGGAACG | | | | |
| Ftx TSS2 F | TGGATCCTAGCCCTTACACG | TSS2 (Ftx) | | 60 | This paper |
| Ftx TSS2 R | CCGGAAATTTGCTCTTTTAGC | | | | |
| Xite A F | ATGGCTTTAAGTCTGTAGCACAA | Xite A | | 60 | (Kung *et al*, 2015) |
| Xite A R | CAGCCTCTATTCAGCTAGACTCC | | | | |
| Xite C F | CAAGGTTGGGAACAAGGTATATCAGG | Xite C | | 60 | (Kung *et al.*, 2015) |
| Xite C R | GGACAAGGGACAGAAGTGCTTATTTTAC | | | | |
| Tsix P2 F | ACTCCCTCAAACCCTCAGTG | 1 (Tsix) | Tsix major promoter | 60 | This paper |
| Tsix P2 R | CACGCTCTTCTTCCATCACG | | | | |
| Dxpas34 F | CGTCGAGGATCTGGGCTATC | 2 (Tsix) | Dxpas34 region | 60 | This paper |
| Dxpas34 R | TAGTGACCTCCCAGTAAGCG | | | | |
| Tsix 33 F | ACGCTTTGCATATCTACCTGTAACA | 3 (Tsix) | Downstream of Dxpas34 | 60 | (Navarro *et al*, 2010) |
| Tsix 33 R | CTGGGCAGAGCAGAGGTGA | | | 60 | |
| Rif1 Peak F | CTGAGTCATGCTGAGGACCG | Peak | Enriched for RIF1 in mESC | 60 | This paper |
| Rif1 Peak R | GCCGGCTCAGCTTCTCC | | | | |
| cRAD F | TAAAGCAGATGGCAGACACG | cRAD | Enriched for RIF1 in mESC and MEF | 60 | This paper |
| cRAD R | TCCGGAAAATACCCAGAGTG | | | | |
| Intergenic1 F | CCAGTAGCAAAAGGGCTCTG | Inter1 | | 60 | This paper |
| Intergenic1 R | CCCAAAATCCCATGTCAAAC | | | | |
| Intergenic2 F | GCAGTGATGTCCACAAGGGA | Inter2 | | 60 | (Karnowski *et al*, 2008) |
| Intergenic2 R | GTGCCTCATGTGCAGTCAGT | | | | |
| Ezr F | GGCCCCGTAACTGCTCTTTA | Ezr | | 60 | (Ding *et al.*, 2018) |
| Ezr R | AGTATAAGACGCTGCGGCAA | | | | |
| Znf629 F | TTGGCTCCTGGTGGCGGAGT | Znf629 | | 60 | (Voon *et al*, 2015) |
| Znf629 R | GCCCGGAGAGCTCAGGGTGA | | | | |
| Peg3 F | CTATGGGGTGCTAGCTCCTC | Peg3 | | 60 | This paper |
| Peg3 R | GACTGAGTGAGGGTCTAGGC | | | | |

#### Supplemental Table 4
RT-qPCR primers

| **Primer** | **Sequence 5' – 3'** | **Description** | **Tm (°C)** | **Reference** |
|---|---|---|---|---|
| Xist ex7 F | TAAGGACTACTTAACGGGCT | Xist Exon7 | 60 | This paper |
| Xist ex7 R | TACTCAGACATTCCCTGGCA | Xist Exon7 | 60 | This paper |
| Xist ex3 F | GGACTGCCAGCAGCCTATAC | Xist Exon3 | 60 | This paper |
| Xist ex4 R | CTCCACCTAGGGATCGTCAA | Xist Exon4 | 60 | This paper |
| Xist RT_129 F | TAGCTGAAGTCTACGCCTCT**G** | *129*-specific Xist Exon7 | 60 | This paper |
| Xist RT_cast F | GTAGCTGAAGTCTACGCCTCT**A** | *castaneus*-specific Xist Exon7 | 60 | This paper |
| Xist RT_129/cast R | CATTCAAGCCCCGTTATGTTC | Xist Exon7 | 60 | This paper |
| Tsix ex4 F | CTGTGAATTATTTGTCAGCGTGAATCA | Tsix Exon4 | 60 | (Shibata *et al*, 2008) |
| Tsix ex4 R | AGAGATCAGACACCCTGGGTATTAG | Tsix Exon4 | 60 | (Shibata *et al.*, 2008) |
| Fgf5 F | AAAGTCAATGGCTCCCACGAA | | 60 | (Poh *et al*, 2014) |

| | | | | |
|---|---|---|---|---|
| Fgf5 R | CTTCAGTCTGTACTTCACTGG | | 60 | (Poh *et al.*, 2014) |
| Oct6 F | CTCAAGCCGCTGCTCAAC | | 60 | (Malaguti *et al*, 2019) |
| Oct6 R | CGCGATCTTGTCCAGGTT | | 60 | (Malaguti *et al.*, 2019) |
| Pax6 F | ACCAGTGTCTACCAGCCAATC | | 60 | (Poh *et al.*, 2014) |
| Pax6 R | GCACGAGTATGAGGAGGTCTGA | | 60 | (Poh *et al.*, 2014) |
| Hand1 F | CACCACCTACCACCGCAGTA | | 60 | (Poh *et al.*, 2014) |
| Hand1 R | CCTTCTTGGGTCCTGAGCCTTT | | 60 | (Poh *et al.*, 2014) |
| Rex1 F | GAGCTGAACTCCTAGCCGCCTAGATT | | 60 | This paper |
| Rex1 R | TTTGGTCAGTGGTATTTGGGGAGA | | 60 | This paper |
| Nanog F | GCATCTTCTGCTTCCTGGCAA | | 60 | This paper |
| Nanog R | GAACTATTCTTGCTTACAAGGGTCTGC | | 60 | This paper |
| Oct3-4 F | GGCGTTCTCTTTGGAAAGGTGTTC | | 60 | (Stavropoulos *et al*, 2001) |
| Oct3-4 R | CTCGAACCACACATCCTTCTCT | | 60 | (Stavropoulos *et al.*, 2001) |
| Sox2 F | GCTCGCAGACCTACATGAAC | | 60 | (Shimoda *et al*, 2007) |
| Sox2 R | GCCTCGGACTTGACCACAG | | 60 | (Shimoda *et al.*, 2007) |
| Gata4 F | CTAACCGGCCCCTCATTAGG | | | (Spruce *et al*, 2010) |
| Gata4 R | CACCCTCGGCATTACGACG | | | (Spruce *et al.*, 2010) |
| Gapdh F | TGTGAGGGAGATGCTCAGTG | | 60 | This paper |
| Gapdh R | ATGGCCTTCCGTGTTCCTAC | | 60 | This paper |
| Cdx2 F | AGGCTGAGCCATGAGGAGTA | | 63.8 | (Shimosato *et al*, 2007) |
| Cdx2 R | CGAGGTCCATAATTCCACTCA | | 63.8 | (Shimosato *et al.*, 2007) |
| Ubiquitin F | GATCCTCTTACCCCCTCGTC | | 60 | (Ruffell *et al*, 2009) |
| Ubiquitin R | CCTTTAGGCCACTCCTTCCT | | 60 | (Ruffell *et al.*, 2009) |
| β-Actin F | AGTGTGACGTTGACATCCGT | | 60 | This paper |
| β-Actin R | TGCTAGGAGCCAGAGCAGTA | | 60 | This paper |
| Rplp0 F | TCCAGAGGCACCATTGAAATT | | 60 | (Furlan *et al*, 2018) |
| Rplp0 R | TCGCTGGCTCCCACCTT | | 60 | (Furlan *et al.*, 2018) |
| Sdha F | GCTCCTGCCTCTGTGGTTGA | | 60 | (Veazey & Golding, 2011) |
| Sdha R | GCAACACCGATGAGCCTG | | 60 | (Veazey & Golding, 2011) |
| 18S F | CATGTCTAAGTACGCACGGC | | 60 | This paper |
| 18S R | CAAGTAGGAGAGGAGCGAGC | | 60 | This paper |

### Supplementary Methods

#### Animal care and use

All mice used in the study were housed and bred in the Animal House located at the EMBL Rome (Epigenetics & Neurobiology Unit). All mice were housed in ventilated cages on a standard dark/light cycle. All procedures involving mice adhered to the guidelines in accordance with European Legislation that exists for the protection of animals used for experimental and other scientific purposes (European convention ETS123/Council of Europe, European directive 86/609/EEC and the more recently published Directive 2010/63/EU and its Italian implementation directive 2014/26) as well as the current Guidelines of International Organizations such as the Association for the Assessment and Accreditation of Laboratory Animal Care International - AAALAC and the Federation of European Laboratory Animal Science Association – FELASA.

#### Gene targeting

The targeting strategy deletes exons 8 and 9, replacing them with a 3-frames STOP cassette, a bovine growth hormone polyadenylation site and a Neomycin-resistance gene. The splicing of exons 6 or 7 to exon 10 generates a frame shift. The targeting vector was generated by recombineering[1], using bacterial artificial chromosome RCPI-23 395O2 (American Type Culture Collection) in *Escherichia coli* strain EL350. The construct was sub-cloned by Xho I digest into the pDTA-TKIII vector, which allows double negative selection (*DTA* and *TK* genes), linearized with NotI, and electroporated into Bruce4 C57BL/6 ES cells. Two independent ES cell clones were injected by standard techniques into 129Svj blastocysts, and chimeras were evaluated based on coat color. Chimeric founders were crossed to C57BL/6J females, and the F1 mice were either maintained in a mixed genetic background by inter-crosses or back-crossed at least 8 times in a pure C57BL/6J background. Genotyping of mice and derived cells was performed by PCR with the following primers: genRif1 comm (r) 5'-caataccagcctcagctacattc-3'; gen Rif1 KO wt (f1) 5'-tgtggagtccatgtcttttgc-3'; and genRif1 KO2 (f2) 5'-atagcctgaagaacgagatcagc-3'. Annealing temperature: 64.3 °C, 40 cycles. PCR generates a 154-bp product for the wild-type allele, a 322-bp product for the deleted (DEL) allele. The southern probes p1 (exon 5) and p2 (exon 11) were generated by PCR from BAC RCPI-23 395O2, with the following primers: p1 oldE1a,

5'-cctcaaataaacctccctta -3; and oldE1s 5'-cagttggatttatagatgtgc-3'; and p2 oldE5a, 5'-ctaatgacagagcaaggtga -3'; and oldE5s 5'-cttttacacacttgacaatg-3'.

**Mouse cell Lines and derivation of embryonic stem cells**

The *Rif1* knockout mouse line was obtained by targeting the *Rif1* locus in Bruce 4 embryonic stem cells. The targeting construct inserts a three-frames STOP cassette and a Neomycin resistance gene, followed by a lox P site into exon 8. Exon 9 is deleted. The targeted cells were screened by PCR and the positive clones verified by Southern blot, following BsrGI and EcoRV digest of the genomic DNA. Two independent clones were injected into blastocysts. Chimeric mice were bred and progeny PCR-genotyped to verify germline transmission. Mice are genotyped using a three-primer PCR Rif1KO comm: 5'-caataccagcctcagctacattc-3'; Rif1 KO wt: 5'-tgtggagtccatgtcttttgc-3'; Rif1 KO2 5'-atagcctgaagaacgagatcagc-3'. Rif1 wild type allele produces a 154 bp band, Rif1 knock-out a 322bp band. $Rif1^{+/-}$ $XY^{-sry}$ $Sry^{Tg}$ mice were obtained by crossing the $XY^{-sry}$ $Tg^{Sry}$ mice[2] to $Rif1^{+/-}$, followed by inter-crosses of the progeny. $Rif1^{F/F}$ $Rosa26^{CreERT/CreERT}$ ($Rs26^{CreERT}$ -*Mus musculus C57BL/6J-129/SvJ*)[3] were crossed with *Mus musculus castaneus* mice and the F1 were inter-crossed for derivation of mESCs of the desired genotypes ($Rif1^{F/F}$ $Rs26^{+/CreERT}$ and $Rif1^{+/+}$ $Rs26^{+/CreERT}$). Derivation of ES cells was carried out according to the protocol as described previously[4-6]. Briefly, Blastocyst outgrowth was cultured on passage 1 Mitomycin C (#M4287, Sigma-Aldrich) -treated primary mouse embryonic fibroblasts (pMEFs=feeders) plated at a density of about 350,000 cells/cm$^2$. First passage disaggregation was performed by mechanical disruption and 0.05% trypsin and the cells re-plated onto feeder coated plates. Once large colonies had formed, they were passaged as ESC lines and expanded in ES medium, Knockout-DMEM (Gibco 10829018), 12.5% heat-inactivated fetal bovine serum (Pan-Biotech), 1% Penicillin/Streptomycin (Gibco 15070063), 1% L-Glutamine (Gibco 25030024), 1% non-essential amino acids (Gibco 11140035), 0.1 mM 2-Mercaptoethanol (Gibco 31350010) and supplemented with 20 ng/ml leukemia inhibitory factor (LIF, EMBL Protein Expression and Purification core facility) and 1 μM MEK inhibitor PD0325901 and 3 μM GSK3 inhibitor CHIR99021 (The University of Dundee, Division of Signal Transduction Therapy)-2i. When cells were stably growing, gelatin adaptation was performed by plating the mESCs on a gelatinised plate (0.1% bovine gelatin in PBS,

#G9391 Sigma-Aldrich) with decreasing numbers of feeders. Cells were cultured at 37 °C in 7.5% $CO_2$. To generate a homogeneous population of cells carrying two X chromosomes, female cell lines with the desired genotype (ESC 34= *Rif1*$^{+/+}$ *Rs26*$^{+/+}$ ; ESC 15 and ESC 16= *Rif1*$^{F/F}$ *Rs26*$^{+/CreERT2}$, ESC 5= *Rif1*$^{+/+}$ *Rs26*$^{+/CreERT2}$) were sub-cloned by plating cells at very low density on gelatinised tissue culture plates. After 3-5 days, single colonies were picked, expanded and characterized. In parallel, the male lines were subcloned and karyotyped too. For knock down studies, the mESC line 34 was used. The Fa2L mESC line[7] used in this study is a subclone (S4) of the Fa2L line (kind gift from P. Avner), adapted to culture in 2i-supplemented medium. The Fa2L line is a female *129/castaneus* hybrid mESC line that carries the insertion of a premature termination site in the *Tsix* gene. As a consequence of this mutation, when these cells are differentiated, the chromosome carrying the mutation (the *129* allele) is almost always selected to become the inactive X chromosome. The Fa2L S4 cell line was maintained on gelatinised plates as described above.

**Whole-mount embryo staining**

Embryos were isolated from *Rif1*$^{+/-}$ inter-crossed timed matings at the appropriate day of gestation, by either flushing in M2 medium (Sigma-Aldrich M7167) or by dissection of the uterus and zona pellucida. Immunostaining for Oct4 and H3K27me 3 was performed as in [8,9]. In brief, by mouth-pipetting, the isolated embryos were transferred into fixative (4% PFA in PBS) for 15 minutes (min.) at RT, into wash buffer (3 mg/ml PVP in PBS) and finally into permeabilization buffer (0.25% Triton X-100, #T9284, Sigma-Aldrich in PBS/PVP) for 30 min. All the procedures were performed in siliconized watch glasses. Blocking was performed for 15 min. in 2% donkey serum, 0.1% BSA, 0.01% Tween-20, in PBS (blocking solution). The primary antibodies were diluted in blocking solution and embryos were incubated in primary antibody for 2 days at 4°C. After 3 washes of 4 hours each in PBS-T, embryos were transferred into secondary antibodies diluted in blocking solution (Molecular Probes Alexa, 1:500) and incubated for 24 hours. After 3 washes of 4 hours each in PBS-T, the embryos were taken through a series of 25, 50, 75, 100% Vectashield (with or without DAPI). The embryos were mounted in a small drop of Vectashield on a slide surrounded by drops of vaseline. A 13x13mm coverslip was gently lowered on to the vaseline drops and pressed lightly to immobilise the embryos. The coverslips were then sealed with nail varnish. After imaging, the coverslips were gently removed and the embryos recovered by mouth pipetting to be transferred into genotyping buffer (100 mM Tris HCl pH 8, 0.5% IGEPAL #I3021, Sigma-Aldrich, 0.5% Tween-20, 0.2 mg/mL proteinase K; #P6656, Sigma-Aldrich) and digested at 55 °C at least 3-4 hours. 1 μL of lysate was used for the first of two rounds of nested PCRs to determine the *Rif1* genotype. The nested PCR was performed with the following primers: PCR 1: genRif1KO comm I: 5'-gattagtttgggctacctacttagttc-3', genRif1 KO wt, 5'-tgtggagtccatgtcttttgc-3', and genRif1 KO2: 5'-atagcctgaagaacgagatcagc-3', to give 180bp wild type and 342 bp KO bands. 3μl of the first PCR were used for PCR 2, with the primers: genRif1WT3: 5'-gagttaaacatctgcaagcctga-3', Rif1KO comm, and genRif1 KO: 5'-ccaattcgccctatagtgagtc-3', that give a 92bp band for the KO and 166bp band for the wild type alleles. For PCR 1, 30 cycles and an annealing temperature of 55.7°C were used; for PCR 2, 25 cycles and the same annealing temperature. The sex of the embryos was assigned by H3K27me3 staining of the extra-embryonic tissues.

##### E7.5 embryos imaging and genotyping

The embryos were first imaged, focusing on one or two, as seen in Fig. 1D. They were then aligned, a picture taken again to document the order and match each embryo to the individual/pair picture, and recovered one by one, in order. The whole embryo was dissolved in genotyping buffer, and the extracted DNA was used for genotyping and sex-typing PCR. The primers used for determining the sex of the embryos are the following: X chr D79F1: 5'-aataaatgttttacaactcctgattcc-3; X chr D79R4: 5'-tgcatagacgtgtaaaacctgc-3'; Y chr SRY1: 5'-gagagcatggagggccat-3'; Y chr SRY2: 5'-ccactcctctgtgacact-3', annealing temperature 60°C. The sex-determining PCR gives a 194bp band for the X chromosome and a 265bp band for the Y chromosome.

##### Luciferase reporter assay

The firefly luciferase reporter plasmids used: pGL4.10(luc2) (control empty Firefly luciferase vector; Promega E6651) and pGL4.10(luc2)-Xist-2p (Xist-2p-luc; Xist reporter construct with Xist promoter and proximal part of exon 1 were kindly donated by Cristina Gontan[10]). For each transfection reaction Lipofectamine 3000 reagent (Thermo Scientific L3000008; 3.75μl per transfection) was used to transfect 2.5 x $10^5$ cells (mESCs of either line 34 or line 16) in 2.5 ml KO DMEM with OHT in a non-coated $3cm^2$ plate after 2 days of OHT treatment at 0 days of EB differentiation (Fig. 2A) with

2.75 μg total DNA comprising 2.5 μg luciferase reporter plasmid (as described above) and 0.25 μg of pRL-SV40 Renilla luciferase plasmid as a transfection control. The luciferase assay was conducted 48 hours after transfection, at 2 days of EBs differentiation, using the Dual-Luciferase Reporter Assay System (Promega E1980) and a GloMax luminometer (Promega).

The ratio of firefly and Renilla fluorescence intensity was determined, and the mean value and standard deviation from three technical repeats and three experiments were calculated.

**shRNA Knockdown Experiment**

To generate shRNA containing plasmids, double-stranded shRNAs were inserted in between the EcoRI and AgeI sites of pLKO.1 (#8453 and #24150, Addgene), third generation lentivirus vectors carrying resistance marker for puromycin or hygromycin selection respectively. For shRNA sequences see Supplementary Table 1. Correct integration was verified by sequencing.

To generate viruses carrying shRNA constructs, 7.4 x $10^6$ HEK239T cells were seeded onto $15cm^2$ tissue culture plates and transfected the following day with lentiviral packaging plasmids: 2.6 µg of pMD-HIV1- Gag/Pol, 2.6 µg pRSV-Rev and 5.2 µg of pMD2.G-VSGV, together with 52 µg of pLKO plasmid carrying the desired shRNA sequence, using $Ca^{2+}$-phosphate transfection method[11]. Medium was changed approximately 16 hours after transfection. The same day of transfection of the packaging cells, 6.5 x$10^6$ mESC were seeded for infection onto $15cm^2$ gelatinised tissue culture plates (for the Fa2L S4 cells: 5.2 x$10^6$ cells were seeded per $15cm^2$ plate). On two subsequent days, starting the day after medium change, viral-containing medium was collected, filtered through a 0.45 µm PES filter, supplemented with Lif and 2i (as described above) and Hexadimethrine bromide (Polybrene, #H9268, Sigma-Aldrich) to a final concentration of 6 μg/ml, and used immediately to transduce the mESCs (days -4 and -3, Fig. 4A). 24 hours following the second transduction (day -2, Fig. 4A) cells were trypsinised and re-plated in medium containing 1μg/ml Puromycin Dihydrochloride (#P8833, Sigma-Aldrich), for selection of cells with stable integration.

For Kap1 knock down in combination with differentiation: the EB differentiation protocol was started at day 2 of puromycin selection (day 0, Fig. 4A). Puromycin was further kept in the medium during the first 24 hours of EB differentiation (day 1, Fig. 4A). Cells were differentiated up to 4 days. For Kap1 knock down and ChIP analysis: puromycin resistant mESCs were collected at day 3 of selection.

For Rif1 knock down in combination with differentiation (Fa2L cells): Fa2L S4 cells were subjected to a second round of infection now using pLKO vectors carrying hygromycin resistance markers. In this second round, mESCs were transduced at day 1 of puromycin selection and day 2 of puromycin selection (=Day 0 EB differentiation). The following day, at day 1 of EB, puromycin was replaced with 180 U/ml of Hygromycin B *Streptomyces sp* (#400051, Calbiochem), which was kept in the medium for the rest of the experiment. Western blotting was performed to verify the knock down throughout the experiment.

**Flavopiridol and triptolide treatments**

Flavopiridol hydrochloride hydrate (#F3055, Sigma-Aldrich) was used to inhibit transcription at a concentration of 500 nM for 4 hours. Triptolide (#T3652, Sigma-Aldrich) was at a concentration of 10 $\mu$M for 1 hour.

**YY1 ChIP**

Cells were collected and crosslinked in 1% formaldehyde in cross-linking buffer (50 mM HEPES pH 7.8, 150 mM NaCl, 1 mM EDTA and 500uM EGTA) for 30 min. at 37 °C. Crosslinking was followed by 5 min. quenching in 0.125 M glycine at RT, washed twice in cold PBS and resuspended in lysis buffer (1% SDS, 10 mM EDTA, 50 mM Tris-HCl pH 8.1, supplemented with protease inhibitor cocktail, #11873580 001, Roche). Chromatin fragmentation was performed using a Bioruptor sonication device (Diagenode) to produce a distribution of fragments enriched between 200 and 300 bp. Immunoprecipitation was performed as described for KAP1 and RIF1 ChIP using $\alpha$-YY1 antibody (see Supplementary Table 2) and IgG only control. ChiP-qPCR primers are presented in Supplementary Table 3.

**Protein extraction, SDS-PAGE and western blotting**

Cells were collected, washed once with cold PBS and resuspended in 2x Laemmli buffer at 10,000 cells/ ul. Samples were homogenised using D Micro-Fine 1ml Insulin Syringe with 29G x 12.7mm Needle (#324891, Becton Dickinson) and boiled for 7-

10min. Proteins were resolved by 5% (RIF1, KAP1 and SMC1) and 10% (RNF12 and LMNB1) SDS-PAGE, transferred to 0.45 μm nitrocellulose membrane (#10600002, Amersham), overnight at 4 °C in transfer buffer (390 mM Glycine, 48 mM Tris base, 0.1% SDS and 20% Methanol) or 3 hours (1x Tris-Glycine buffer supplemented with 20% methanol) using standard procedures. Following transfer, membranes were blocked with 5 % (w/v) skim milk powder (#84615.0500, VWR) in 0.05% (v/v) Tween-TBS (TBS-T) for a minimum of 30 min. at RT. Membranes were incubated either overnight at 4 °C or 3 hours at RT with antibodies (see Supplementary Table 2) diluted in blocking buffer or 2.5% milk/ TBS-T. Secondary antibodies were diluted 1:15,000 in 2.5% milk/ TBS-T and incubated for approximately 45 min. Secondary antibodies were as follow: Donkey Anti-Rabbit IgG Antibody IRDye:800CW Conjugated (#926-32213, LI-COR Biosciences), Donkey Anti-Rabbit IgG Antibody IRDye:680RD Conjugated (#925-68073, LI-COR Biosciences). Goat Anti-Mouse IgG Antibody IRDye:800CW Conjugated (#925-32210, LI-COR Biosciences). Blots were imaged with Odyssey® CLx Imager (LI-COR Biosciences) and the intensity of each band was quantified using the Odyssey imaging software.

**Nuclear-Cytoplasmic Fractionation**

Cell fractionation and solubilisation of chromatin-bound proteins by salt extraction was performed according to[13]. Briefly, mESCs were harvested, washed 2x in PBS and resuspended in cold Buffer A (10mM HEPES pH 7.9, 10mM KCl, 1.5mM $MgCl_2$, 0.34M sucrose, 10% glycerol, 1mM DTT with freshly added Protease (#11873580 001, Roche) and Phosphatase Inhibitors) at 4 x $10^7$ cells/µl making sure no cell clumps remain. Triton X-100 to a final concentration of 0.1% was added slowly, swirling the pipette tip from bottom to top. Samples were incubated 5 min. on ice followed by a centrifugation for 4 min. at 1300 g, 4 °C. The supernatant, containing the cytosolic fraction (S1) was collected and spun in an ultracentrifuge at 20000g for 15 min. and the S1-pellet was resuspended in 2x Laemmli buffer. The remaining nuclear pellet (P1) was gently washed in 500µL Buffer A and centrifuged 4 min. at 1300 g, 4 °C. The washed P1-pellet was resuspended in a volume of Buffer B (3mM EDTA, 0.2mM EGTA, 1mM DTT with freshly added Protease and Phosphatase Inhibitors) equal to the initial Buffer A and incubated on ice for 30 min. Incubation was followed by a centrifugation for 4 min. at 1700 g, 4 °C. The supernatant, containing the soluble nucleoplasmic fraction (P2) was collected, spun down and resuspended in 2x Laemmli buffer. The pellet, containing the nuclear pellet+chromatin fraction (P3) was resuspended in a volume of 2x Laemmli buffer equal to the initial buffer A volume. All samples were syringed through a D Micro-Fine 1ml Insulin Syringe with 29G x 12.7mm needle and boiled for 7-10 min.

**RNA-seq library preparation and analysis**

Two wild type female cell lines (*Rif1*$^{+/+}$ *Rs26*$^{+/\ CreERT2}$) and two female *Rif1* conditional knock-out cell lines (*Rif1*$^{F/F}$ *Rs26*$^{+/CreERT2}$) were used for RNA sequencing analysis. Cell differentiation was combined with Rif1 deletion as described above. Samples were collected before OHT treatment (pre-samples), at day 2 of OHT treatment (ESC-samples) and at 2d EB differentiation (2d EB) samples. The pre-samples were used to normalise for clonal variably. RNA extraction was performed as described above and samples were stored at −80 °C until further processed. Total RNA was assessed using stranded rRNAminus RNA-Sequencing. The GeneCore facility (EMBL Heidelberg) prepared the sequencing libraries and performed the RNA sequencing. The barcoded libraries were pooled and run on an Illumina NextSeq500 sequencer using a single-end mode with a read length of 75 bases.

**Immunofluorescence**

mESCs were plated on gelatinised coverslips overnight. For differentiated cells, at the day of harvest, the embryoid bodies were gently disrupted into single cells using 0.05% Trypsin-EDTA and mechanical dissociation, then cytospun onto polysine coated slides (#J2800AMNZ, Thermo Scientific) at 1000 rpm for 5 min. Cells were fixed in 4% paraformaldehyde in PBS for 10 min. at RT and stored at 4 °C in PBS. For staining, coverslips were permeabilized in 0.5% Triton X-100/ PBS at RT for 10 min., washed x2 with PBS/ 0.2% Tween-20 and blocked for 30 min. in 1% BSA/ PBS/ 0.02% Tween-20. Primary antibody (Supplementary Table 2) was diluted in 0.5% BSA/ PBS/ 0.02% Tween-20 and coverslips were incubated for 3 hours at RT. After x3 washes in PBS/ 0.02% Tween-20, coverslips were incubated with secondary antibody, Alexa-488 Donkey anti-Rabbit antibody (#A21206, Thermo Fisher) diluted 1:800 in 0.5% BSA/ PBS/ 0.2% Tween-20, for 45 min. at RT in the dark. Following x3 washes in PBS/ 0.02% Tween-20 and x1 wash in PBS, coverslips were mounted with Vectashield including DAPI (#H-1200, Vector Laboratories). Images were acquired using a Zeiss Axio Imager. Cell count minimum of 100 cells /slide.

#### RNA FISH

RNA FISH was done as previously described[15]. For the probes preparation, 1µg Xist cDNA was labelled with Green/Red-dUTP (Abbott Molecular) using a nick translation kit (Abbott Molecular) in a total volume of 50 µL. For the Tsix probe, 1 µg WI1-1112M11 fosmid was labelled using Red/Green-dUTP (Abbott Molecular) using a nick translation kit (Abbott Molecular) in a total volume of 50 µL. For every 22x22mm coverslip, 5.5 µL of labelled probes (2.5 µl Xist + 2.5 µl Tsix) were NaCl/ethanol precipitated with 1 µg of Salmon Sperm DNA (Life Technologies), 0.3 µl yeast tRNA (20mg/ml) and 3 µg of mouse Cot-1 (LifeTechnologies), and resuspended in 6 µL of formamide, denatured for 5 min at 75 °C and kept on ice for 5 min. 6 µl of 2X hybridization buffer containing RNAse inhibitors (VRC) was then added to the probes (final concentration: 50% formamide, 2X SSC, 10% dextran sulfate, 1 mg/mL BSA).

For the slides preparation, about 60,000 cells per slides were prefixed on ice (1% formaldehyde) for 10 min., washed in PBS and cyto-spun onto coverslips as previously described (1,800 rpm for 3 min.)[16]. Slides were permeabilised with 0.5% Triton X-100 in PBS + 1% VRC for 5 min. on ice, then washed with PBS and fixed with 4% formaldehyde in PBS for 5 min. on ice. Following fixation, slides were dehydrated with 70%-80%-95%-100% ethanol, and hybridized overnight at 37 °C in humid chamber. The following day, the slides were washed 3 times with 50% formamide, 2X SSC at 42°C, and 3 times with 2X SSC at 42°C. Slides were mounted in Vectashield with DAPI and visualized using a 63x oil immersion objective with the Zeiss Axio Observer Z1 microscope.

#### Motif Detection

Identification and enrichment of known and de novo sequence motifs within Rif1 ChIP-seq peaks was performed with HOMER[17], using the “findMotifsGenome.pl” command. The “-size given” option was used to utilise the full length of each peak, rather than the default option of 200 bp. Motif discovery was performed independently for peaks identified in two biological ChIP-seq replicates. Probability plots were generated using the “ggseqlogo” package in R.

#### Methods references


1 Lee, E. C. *et al.* A highly efficient Escherichia coli-based chromosome engineering system adapted for recombinogenic targeting and subcloning of BAC DNA. *Genomics* **73**, 56-65 (2001).

2 De Vries, G. J. *et al.* A model system for study of sex chromosome effects on sexually dimorphic neural and behavioral traits. *J Neurosci* **22**, 9005-9014 (2002).

3 Buonomo, S. B., Wu, Y., Ferguson, D. & de Lange, T. Mammalian Rif1 contributes to replication stress survival and homology-directed repair. *J Cell Biol* **187**, 385-398, doi:10.1083/jcb.200902039 (2009).

4 Foti, R. *et al.* Nuclear Architecture Organized by Rif1 Underpins the Replication-Timing Program. *Mol Cell* **61**, 260-273, doi:10.1016/j.molcel.2015.12.001 (2016).

5 Bryja, V., Bonilla, S. & Arenas, E. Derivation of mouse embryonic stem cells. *Nat Protoc* **1**, 2082-2087, doi:10.1038/nprot.2006.355 (2006).

6 Bryja, V. *et al.* An efficient method for the derivation of mouse embryonic stem cells. *Stem Cells* **24**, 844-849, doi:10.1634/stemcells.2005-0444 (2006).

7 Luikenhuis, S., Wutz, A. & Jaenisch, R. Antisense transcription through the Xist locus mediates Tsix function in embryonic stem cells. *Mol Cell Biol* **21**, 8512-8520, doi:10.1128/MCB.21.24.8512-8520.2001 (2001).

8 Ohinata, Y., Sano, M., Shigeta, M., Yamanaka, K. & Saitou, M. A comprehensive, non-invasive visualization of primordial germ cell development in mice by the Prdm1-mVenus and Dppa3-ECFP double transgenic reporter. *Reproduction* **136**, 503-514, doi:10.1530/REP-08-0053 (2008).

9 Seki, Y. *et al.* Extensive and orderly reprogramming of genome-wide chromatin modifications associated with specification and early development of germ cells in mice. *Dev Biol* **278**, 440-458, doi:10.1016/j.ydbio.2004.11.025 (2005).

10 Gontan, C. *et al.* RNF12 initiates X-chromosome inactivation by targeting REX1 for degradation. *Nature* **485**, 386-390, doi:10.1038/nature11070 (2012).

11 Graham, F. L. & van der Eb, A. J. A new technique for the assay of infectivity of human adenovirus 5 DNA. *Virology* **52**, 456-467, doi:10.1016/0042-6822(73)90341-3 (1973).

12 Bulut-Karslioglu, A. *et al.* A transcription factor-based mechanism for mouse heterochromatin formation. *Nat Struct Mol Biol* **19**, 1023-1030, doi:10.1038/nsmb.2382 (2012).

13 Mendez, J. & Stillman, B. Chromatin association of human origin recognition complex, cdc6, and minichromosome maintenance proteins during the cell cycle: assembly of prereplication complexes in late mitosis. *Mol Cell Biol* **20**, 8602-8612, doi:10.1128/mcb.20.22.8602-8612.2000 (2000).

14 Livak, K. J. & Schmittgen, T. D. Analysis of relative gene expression data using real-time quantitative PCR and the 2(-Delta Delta C(T)) Method. *Methods* **25**, 402-408, doi:10.1006/meth.2001.1262 (2001).

15 Moindrot, B. *et al.* A Pooled shRNA Screen Identifies Rbm15, Spen, and Wtap as Factors Required for Xist RNA-Mediated Silencing. *Cell reports* **12**, 562-572, doi:10.1016/j.celrep.2015.06.053 (2015).

16 Cerase, A. *et al.* Spatial separation of Xist RNA and polycomb proteins revealed by superresolution microscopy. *Proceedings of the National Academy of Sciences of the United States of America* **111**, 2235-2240, doi:10.1073/pnas.1312951111 (2014).

17 Heinz, S. *et al.* Simple combinations of lineage-determining transcription factors prime cis-regulatory elements required for macrophage and B cell identities. *Molecular cell* **38**, 576-589, doi:10.1016/j.molcel.2010.05.004 (2010).
